# Insights into the conformational dynamics of the cytoplasmic domain of metal-sensing sensor histidine kinase ZraS

**DOI:** 10.1101/2024.11.23.624959

**Authors:** Nilima Mahapatra, Pranjal Mahanta, Shubhant Pandey, Rudresh Acharya

**Affiliations:** School of Biological Sciences, National Institute of Science Education and Research, Bhubaneswar 752050, Odisha, India; Homi Bhabha National Institute, BARC Training School Complex, Anushakti Nagar, Mumbai 400094, Maharashtra, India

**Keywords:** ZraS, sensor histidine kinase, two-component system, cytoplasmic domain, trans-autophosphorylation, *Escherichia coli*

## Abstract

ZraS is a metal sensor integral to ZraPSR, a two-component signaling system found in enterobacter*s*. It belongs to a family of bifunctional sensor histidine kinases (SHKs) and is speculated to sense zinc-induced stress on the bacterial envelope. Information on the structure-function relationship of sensor kinases is elusive due to the lack of full-length structures, intrinsically dynamic behavior, and difficulty trapping them in active state conformations. While the kinase domains of a few sensor histidine kinases are well-characterized, they exhibit significant functional diversity attributed to their modular multi-domain arrangement in the cytoplasmic region, combined with other signal transducing elements such as simple helices, HAMP and PAS domains. We report the crystal structure of the entire cytoplasmic region of *Escherichia coli* ZraS (EcZraS-CD) resolved at resolution 2.49 Å, comprising of a unique helical linker and the kinase domain. In the asymmetric unit, four molecules of ZraS assemble as homodimers trapped as two ligand-bound occluded conformers. Our analysis using these conformers show that modulation of the dimer bundle through segmental helical bending, sliding, and rotation leads to reorganization of the dimerization interface during kinase activation. Further, our analysis reveals the significance of aromatic amino acid interactions and loop residues at the dimer base in regulating the directionality of rotation during autophosphorylation. We also performed an invitro coupled assay to determine ATPase activity. Overall, our findings provide structure-based mechanistic insights into the process of autophosphorylation in trans-acting sensor histidine kinases.

## Introduction

Bacterial cells thrive in rapidly fluctuating environments, requiring continual detection and conditional adaptation. These cells have an intricate network of biosensors that regulate sense-response cascades, aiding in their protection, persistence, and proliferation. Fundamental signal transduction events are triggered by environmental cues (Figure 1A) that relay signals into the internal cellular machinery, thereby reprograming it to perform essential functions and evade unfavorable conditions. A repertoire of kinases at the cell interface undergoes reversible phosphorylation of conserved histidine residues upon cue detection ^1^. Sensor histidine kinases (SHKs) or histidine protein kinases (HPKs) represent the sensor modules of the sensor-effector dual module systems known as two-component systems (TCSs). They are coupled with cognate response regulators (RRs) eliciting effector functions, generally acting as transcription factors ^2^. Ubiquitously distributed across all three domains (eubacteria, archaea, and eukarya), the implication of sensor kinases in the directing physiological, biochemical, and genetic pathways is remarkable ^1,3^. Absence in humans and their role in regulating antibiotic resistance and virulence make them potential targets for developing novel antimicrobial agents ^4^.

**Figure 1.**
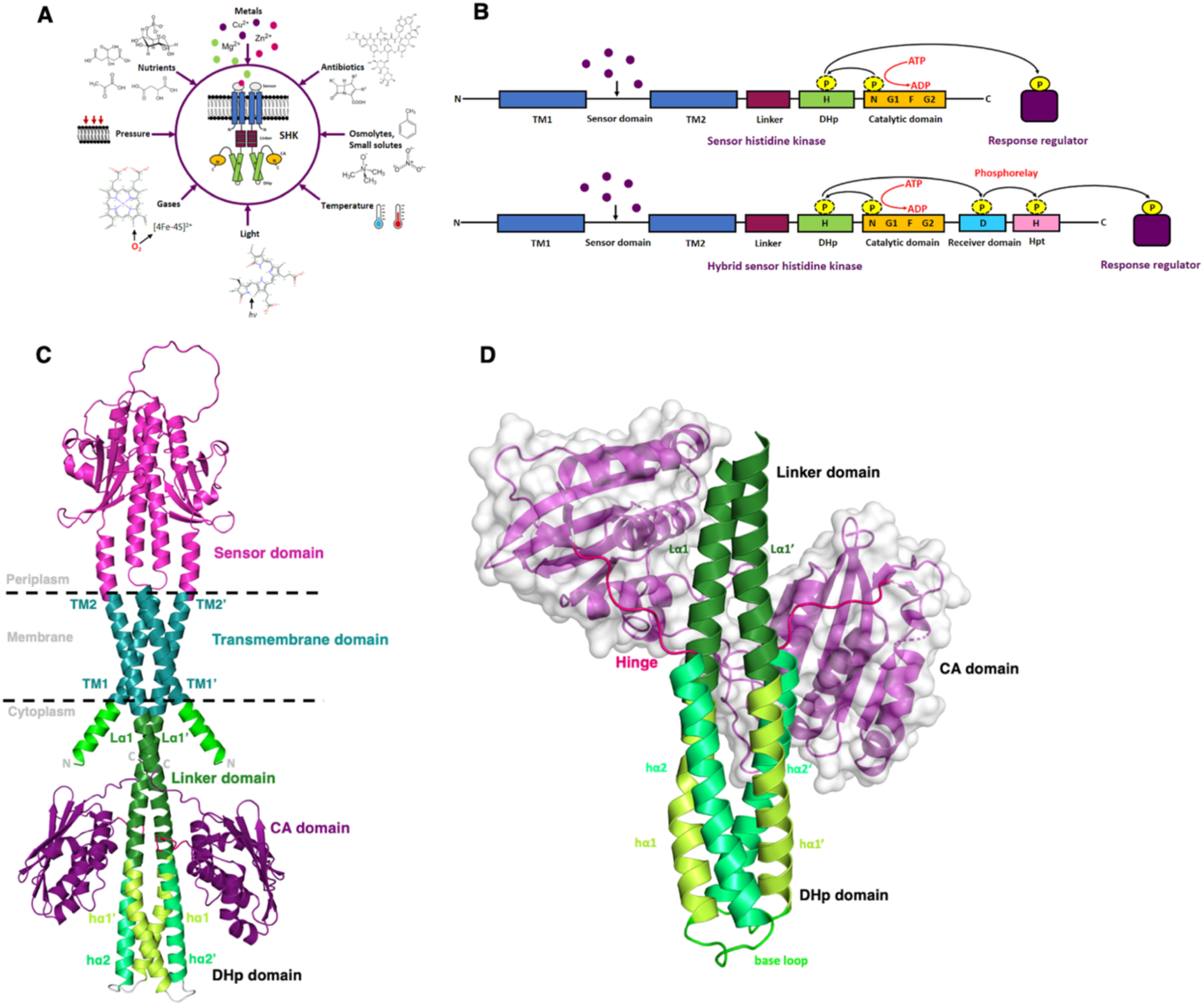
Full-length model and truncated crystal structure of cytoplasmic domain of EcZraS. (A) Schematic representation of different types of stimuli detected by SHKs. (B) Representation of a prototypical SHK showing direct phosphate transfer and a hybrid kinase transferring phosphate through a relay mechanism involving other intermediate domains such as Hpt and receiver domains. (C) AlphaFold model of the apo form of full-length ZraS (EcZraS-F) displaying domain distributions across periplasmic (light magenta), transmembrane (deep teal), and cytoplasmic regions (forest, limon, lime green, and deep purple) ^30^. (D) Three-dimensional crystal structure of ligand-bound ZraS (EcZraS-CD) trapped in an intermediate state. The overall structure of the cytoplasmic region has a simple linker domain (forest) and a kinase domain made of the DHp and CA domains. The DHp region (limon and lime green) is connected to the CA domains (deep purple) using hinges (hot pink).

Prototypical SHKs are homodimers comprising multiple domains distributed across periplasmic, transmembrane, and cytoplasmic regions. The extra-cytoplasmic region typically senses the cue, with two or more transmembrane helices connecting the extra-cytoplasmic domain to the cytoplasmic domain. The cytoplasmic region has multiple domains including the kinase module, possessing one or more domains involved in phosphate acquisition and transfer ^5^. Initiation of signal transduction occurs through self-phosphorylation at a conserved His residue in the central bundle of the cytoplasmic kinase domain. Following autophosphorylation, the transfer of phosphate from His to a conserved Asp residue in RRs activates the cognate RRs. Phosphate acquisition occurs through the catalytic domain (CA) and is transferred to a single dimerization and histidine phosphoacceptor domain (DHp) or relayed through multiple cytoplasmic domains like histidine-containing phosphotransfer domain (Hpt) and receiver domain before reaching the response regulator (Figure 1B and S1A). Hydrolysis of phosphate bound to Asp in RRs post-transcriptional regulation terminates the cascade. An ancillary phosphatase activity in bifunctional SHKs supports phosphate removal from RRs ^6^. It is particularly intriguing to discern the mechanism behind the kinase and phosphatase activities in SHKs.

The current knowledge on signal transfer and integration is limited to the structural and functional characterization of a few sensor kinases ^5^. Structural elucidation of SHKs in different signaling stages, including autophosphorylation, phosphate transfer, and dephosphorylation, is of significance in comprehending the molecular basis underlying the processes. In the absence of full-length structures, evidence on conformational dynamics has been acquired from studies on truncated cytoplasmic stretches. Studies on functional aspects of the cytoplasmic stretch of VicK with essential domain combinations demonstrated enzymatic activity comparable to its full-length kinase ^7^. Isolated kinase domains of DosS and DosT retained their autophosphorylation and phosphate transfer activity ^8^. The osmosensing ability of native EnvZ was also found to be unaltered in the absence of transmembrane segments, making truncated domains appropriate study alternatives ^9^. Nucleotide-bound structures of truncated cytoplasmic regions of CpxA, native and chimeric EnvZ, DesK, CovS/VicK-like, WalK/VicK, YF1, and HK853 determined by X-ray crystallographic studies illustrated multiple conformational states, having diverse intermediate linker domains ranging from simple helices to complicated tertiary folds ^10–18^. Regulation of catalysis through state transitions supported by helical bending and rotation, coordinated with moveable ATP-binding CA domains was reported for DesK. The positioning and orientation of the CA domains in a kinase-active state followed an asymmetric disposition as opposed to phosphatase-active state having a rather symmetric disposition ^12,19^. An asymmetric dimer arrangement facilitated through tilting and rotation of linker domain helices in PhoQ allowed dissociation of an inactive CA domain-central bundle interaction, to attain a kinase-active conformation^20^. EnvZ and VicK also confirmed asymmetric bending of central bundle helices, allowing sequential binding of ATP-binding domains, facilitating self-phosphorylation of alternate His residues in the bundle ^11,13^. Molecular dynamics simulations on the kinase domain of WalK proposed helical bending in the actively phosphorylating monomer to be an indirect process supported by intermediate states having differential kinase bundle helix angles ^14^. Combination of superhelical axis rotation, bending and local unwinding in the kinase core was observed during autophosphorylation of the light sensor YF1, having a simple α-helical linker ^16,21^. Identification of residues involved in autophosphorylation activity were identified through mutational analysis in kinases such as PhoQ, VicK and HK853 ^13,15,18,20^.

Further, the complexity of SHKs is marked by their classification into 11 subfamilies (HPK 1-11), based on conserved kinase regions by Grebe and Stock (1999) and into Type I-V, based on phylogenetic studies by Kim and Frost (2001) ^22,23^. High intrinsic flexibility and instability in-vitro pose a challenge towards isolation of these enzymes for study. Adding to this intricacy, the cytoplasmic stretches have a modular architecture with one or many intermediate domains forwarding the input from transmembrane helices to the kinase domain. Another requisite in a multi-module signal transfer is to determine the contribution of individual modules towards the process, alone and in combination with other signal-transduction elements. Structural reorganization of intermediate linker domains such as HAMP in PhoQ and PAS (Per-ARNT-SIM) in ThkA were found to influence the kinase-phosphatase conformational switching ^24,25^. Presence of HAMP and PAS domains favored a phosphatase-active state, while the enzymes were intrinsically inclined towards a kinase-active state. Considering a handful of known representatives from SHK sub-families detecting multiple stimuli, differing in sequence composition, domain organization, and mode of SHK activation, an understanding of the sequence of events is insufficient. A combination of domains correlated with differential global behavior demands the study of other members of SHK sub-families to understand the mechanism of signal transduction.

Adding to the existing information, here we have characterized the cytoplasmic stretch of a sensor histidine kinase ZraS from *Escherichia coli* to deduce the mechanism of SHK signaling using a simple α-helical signal transducing linker element. ZraS is a membrane-bound stress induced SHK of HPK4 or Type IC orthodox kinase subfamily encoded by an operon zraPSR, functionally homologous to CpxPAR ^22,23,26^. Involvement of zraPSR in preservation of membrane integrity and their absence resulting in susceptibility towards antibiotics in *E. coli* is substantial ^27^. ZraS is also known to be activated upon binding metals such as zinc and lead, released into the periplasmic space upon disruption of the bacterial membrane during envelope stress^28^ (Figure S1B). Metals play a pivotal role as co-factors for metalloproteins and inorganic catalysts that control basic biological processes. Thus, maintenance of metal homeostasis through the detection, uptake, removal, or sequestration of metals is crucial to evade toxicity and facilitate utility. The putative metal binding site is located at a α/β periplasmic domain, that stimulates the cytoplasmic kinase domain through signal transfer via transmembrane and linker helices (Figure 2B) ^29^.

**Figure 2.**
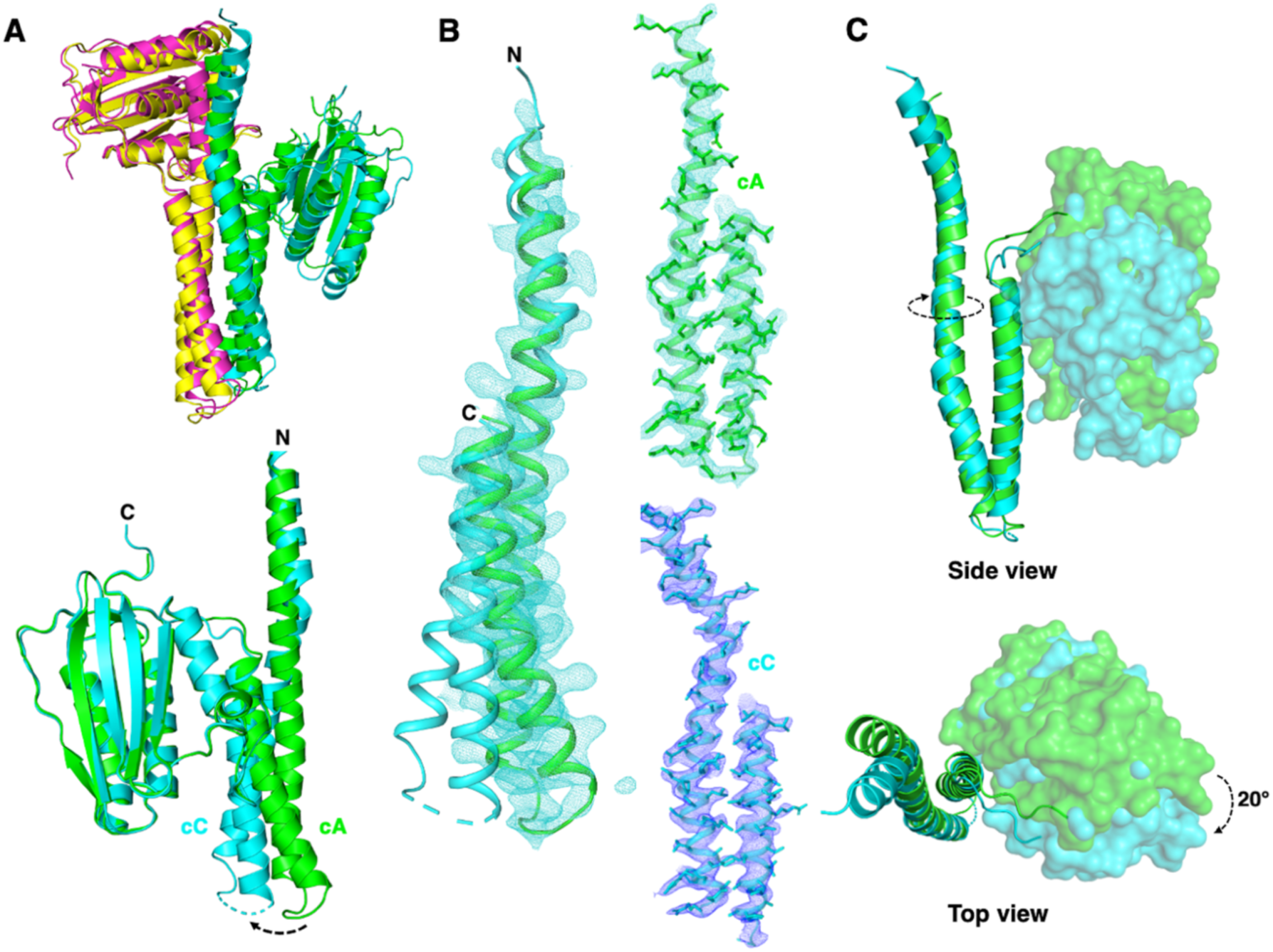
Structural comparison between chains cA of dimer AB and cC of dimer CD. (A) Cartoon representation of the superposed dimers of EcZraS-CD. Chain cC of dimer CD displays a prominent helical bend at the lower region of the DHp bundle compared to chain cA of dimer AB. (B) 2Fo-Fc map contoured at 1.0 σ for the linker and DHp helices of cA and cC. (C) Structural overlap of cA and cC displaying rotation of CA domain through an angle of nearly 20°.

Despite numerous biophysical and biochemical studies on the SHK, the cytoplasmic region remains uncharacterized. According to our findings, the cytoplasmic region of ZraS form functional homodimers. The homodimers trapped during crystallization were found to be in two successive occluded conformations, resembling intermediate kinase-active states. Comparison of these conformers provide structure-aided mechanistic insights into the process of trans-autophosphorylation. We also estimated the rate of ATPase activity for the region using real-time biochemical kinetics.

### Cloning and overexpression

The cytoplasmic domain of metal-sensing protein ZraS from *Escherichia coli* ranging Tyr225-Gly465 (EcZraS-CD) was amplified from its full-length gene template using Q5 high fidelity DNA polymerase with AAAAAACCATGGCGTATCTGCGCTCGCGCC and TTTTTTGGATCCTCATCCTTGTGGGTCCTTACGC as forward and reverse primers having an annealing temperature of 72°C. It was cloned into the multiple cloning site of vector pNIC28-Bsa4 using restriction enzymes NcoI and BamHI and subsequently transformed into *Escherichia coli* DH5α. *Escherichia coli* LEMO21 (DE3) was used for protein production by induction in Luria-Bertani broth, Miller at 0.5 mM isopropyl β-d-1-thiogalactopyranoside after OD 600 reached 0.8 and allowed to grow overnight at 18°C. The clone was designed to produce a 6x-His-tagged protein in the N-terminus with a Tobacco Etch Virus (TEV) protease cleavage site between the affinity tag and the protein.

### Protein purification

The bacterial pellet obtained post-induction was resuspended in lysis buffer (100 mM Tris pH 8.0, 500 mM NaCl, 10% Glycerol, and 20 mM Imidazole) supplemented with 1mM protease inhibitor phenylmethylsulfonyl fluoride (PMSF). Cells were lysed using an ultrasonic processor and centrifuged at 14000 rpm (23,447 xg) for 1 hour at 4°C to remove cell debris. Purification of protein was carried out using affinity chromatography with a HisTrap-FF column. The column was pre-equilibrated using buffer A (25 mM Tris pH 8.0, 200 mM NaCl, 5% glycerol, and 20 mM Imidazole), and the supernatant was allowed to bind. The column was washed with 20 column volume buffer A to remove impurities. Protein eluted at 200-250 mM Imidazole concentration with step and linear gradient elutions of buffer A (25 mM Tris pH 8.0, 200 mM NaCl, 5% glycerol, and 20 mM Imidazole) and buffer B (25 mM Tris pH 8.0, 200 mM NaCl, 5% glycerol and 1M Imidazole). Desalting was performed to remove imidazole from eluted protein by passing it through the HiPrep 26/10 desalting column in buffer C (50 mM Tris pH 8.0, 200 mM NaCl, 2 mM 2-Mercaptoethanol). Desalted protein was incubated with TEV protease overnight at 4°C to remove the 6x-His tag ^31^. TEV protease and uncleaved protein from the mixture were removed by passing the incubated mixture through HisTrap-FF. To get a pure fraction of the desired protein, size exclusion chromatography was performed using Hi-Load 16/600 Superdex 75pg in buffer D (20 mM Tris pH 8.0 and 150 mM NaCl). Concentrated protein was used for structural and biochemical studies.

### Crystallization

EcZraS-CD was concentrated to 8-10 mg/ml and incubated with 1 mM ATP and 5 mM MgCl2. Incubated protein was used to screen the crystallization in commercially available conditions. Crystals were obtained in a condition containing 0.2 M Ammonium sulfate, 0.1 M Tris pH 8.5, and 25% PEG3350 which were optimized using additional reagents to obtain crystals of diffraction quality.

### X-ray diffraction data collection, structure solution, and refinement

Crystals flash frozen with 20% ethylene glycol as a cryoprotectant were used for diffraction at the European Synchrotron Radiation Facility (ESRF). The diffraction datasets were collected at beamline ID30A and processed using *XDS* ^32^. We collected the diffraction datasets for multiple crystals; the datasets were merged and scaled using *XSCALE* ^32,33^. The diffraction was recorded to a maximum resolution of 2.49 Å. The structure was determined by the molecular replacement method implemented in *Phaser* ^34^. For the molecular replacement, we used a polyAla model of the CA domain as a probe, a part of the cytoplasmic kinase domain structure generated by Alphafold^30^. The model was built using iterative refinement (*Phenix*) and tracing the sidechains in the electron density map *(Coot*). Refinements were carried out using *Phenix* ^35^. The electron density map tracing and model building were done using *Coot* ^36^. Data collection and refinement statistics are mentioned in Table-1.

**Table-1.**
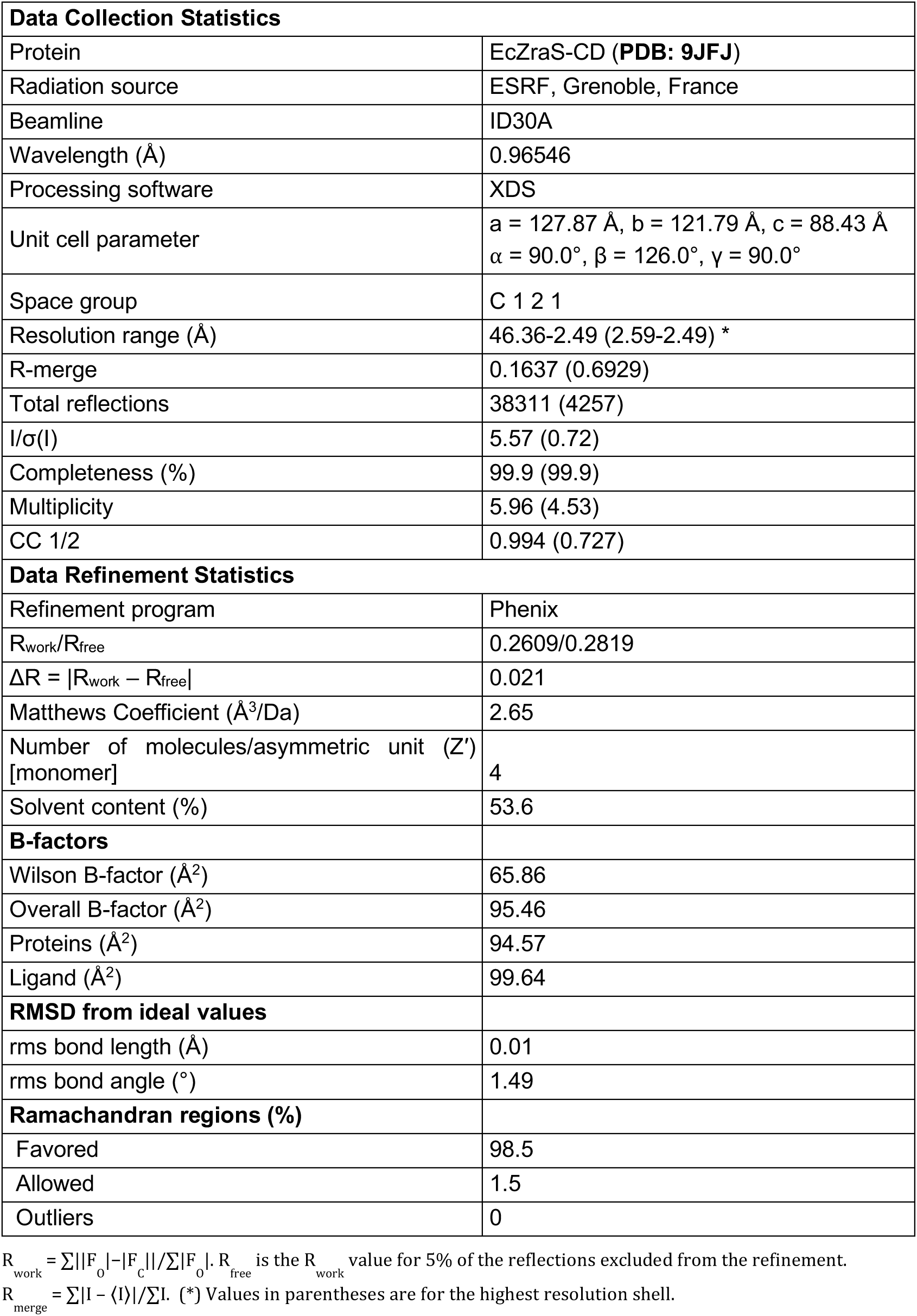
Data collection and refinement statistics of EcZraS-CD.

### Enzyme kinetics

Autophosphorylation and phosphatase activity of the kinase EcZraS-CD was assessed using a continuous pyruvate kinase-lactate dehydrogenase coupled enzyme assay. ATP hydrolysis to ADP is coupled with the oxidation of NADH to NAD^+^ by pyruvate kinase (PK) and lactate dehydrogenase (LDH) in the presence of 2-phosphoenolpyruvate (PEP). PK catalyzes the conversion of PEP to pyruvate, which is reduced to lactate by LDH, catalyzing the oxidation of NADH. We can monitor the NADH depletion, as NADH absorbs at 340 nm (NAD^+^ has no absorbance). A kinetic study was carried out with 15uM protein in a reaction buffer containing 50 mM HEPES pH 8.0, 145 mM KCH_3_CO_2_, 8 mM MgCl_2_, 2 mM phosphoenolpyruvate, 0.16 mM NADH, 4.8 units of pyruvate kinase, 8.1 units of lactate dehydrogenase, with varying ATP concentrations at 25 °C. All reaction components except protein were pre-incubated with ATP at the desired temperature to convert contaminated ADP to ATP in the stock. Protein was then added to start the reaction and decrease in absorbance at 340 nm measured at intervals of 75 seconds up to 20 minutes. The rate of ATP hydrolysis was calculated using linear regression analysis of absorbance values and then fitted using non-linear regression analysis (Michaelis-Menten functions) implemented in GraphPad Prism 5 to determine V_max_ and K_m_ values.

### Sequence and structure-based analysis

Sequence alignment was performed using Clustal-W with representative members of HPK families ^37^. Sequence logos of conserved boxes were generated using Web logo 3.7.12 ^38^. Structural superposition was performed using Superpose and Pair fit algorithms in PyMol ^39^ .Measurement of distances and angles were also made using PyMol. Images for alignments were generated using Boxshade and ESPript 3.0 ^40^. Domain movements were analyzed using PyMol ^39^ and Dyndom ^41^. TWISTER ^42^ and SamCC Turbo ^43^ was used to analyze the coiled-coil regions in the cytoplasmic domain.

### Electrostatic surface charge analysis

Parameters such as atomic charges and radii required to calculate biomolecular electrostatics and solvation were obtained using PDB2PQR ^44^. PQR outputs were used to generate electrostatic potential surface maps in APBS (Adaptive Poisson-Boltzmann Solver) plugin of PyMol with electrostatic potential ranging from -5.0 (red/electronegative) to +5.0 kT/e (blue/electropositive) ^44,45^.

## Results and discussion

### Overall structure of the cytoplasmic stretch of ZraS (EcZraS-CD)

The three-dimensional structure of the truncated cytoplasmic stretch of *Escherichia coli* ZraS (EcZraS-CD) spanning residues Tyr225-Thr459 was solved using molecular replacement and refined to a resolution of 2.49 Å (Table-1). Four molecules of EcZraS-CD denoted as chains cA, cB, cC and cD were traced in the asymmetric unit, assembled as two constitutively parallel homodimers, AB and CD. Doubly-ADP bound states were observed with one ADP and Mg^2+^ ion coordinated with each protomer due to hydrolysis of ATP added during crystallization. The dimers adopt a topology of a classical cytoplasmic domain featuring a linker region: Tyr225-Ala250 and a kinase domain (KD): Gly251-Thr459 (Figure 1D). The kinase domain comprises of a dimerization and histidine phosphoacceptor domain (DHp) and two catalytic and ATP-binding domains (CAs).

The linker helices [Lα1 (cA/cC)/Lα1’ (cB/cD)] from both protomers (Figure S2C) of a homodimer organize into a parallel helical bundle, linking the C-terminus of transmembrane helices (TM2 and TM2’) with the first helix of the kinase domain forming a continuous helical backbone. DHp region is a four-helix bundle made of antiparallel helices, helix-1 [hα1 (cA/cC)/ hα1’ (cB/cD)] and helix-2 [hα2 (cA/cC) / hα2’ (cB/cD)], connected by base loops forming two helix-turn-helix motifs in a homodimer (Figure 1B). The DHp helix-1 (hα1/ hα1’) has the phosphorylable histidine (His254) found in sensor histidine kinases. The addition of γ-phosphate to His254 catalyzes autophosphorylation in ZraS. Contiguous with the helix-2 are hinge regions (Val305-Leu312) connecting the DHp to CA domains. The CA domain is a globular domain with α/β topology comprising of four alpha helices (α1-α4) and seven beta sheets (β1-β7), out of which three alpha helices (α1, α2, and α3) and five antiparallel beta sheets (β2, β4, β5, β6, and β7) are arranged in two layers forming a sandwich-like fold (Figure S2E). A groove formed towards the edge of this sandwich forms the ligand-binding pocket and accommodates the nucleotide. The DHp and CA regions of EcZraS-CD comprise of highly conserved regions found across members of the GHKL (Gyrases, Hsp90, Histidine kinases, MutL) superfamily. Based on single-letter codes of conserved residues and sequence consensus, the regions are named as H (His), N (Asn), G1 (Gly), F (Phe), G2 (Gly), and G3 (Gly) boxes ^15,46,47^. H-box is present in the upper segment of the DHp helices hα1/ hα1’ (Figure S2A). N-box, G1-box, F-box, G2-box, and G3-box are found in the CA domain lining the ligand-binding pocket (Figure S2B). Multiple sequence alignment of the conserved boxes classified ZraS as a member of the HPK4 subfamily similar to NtrB and FleS (Figure S3A). The residues Ala253 and Gln258 precedes the conserved residues His254 and Pro259 respectively in the H-box, a characteristic feature of the HPK4 family of kinases ^22^. A second identifying feature is the TTK motif formed by Thr414-Thr415-Lys416 adjacent to Phe409 of the F-box (Figure S2D and S3B).

## Methods

### Structural comparison of dimers of EcZraS-CD

Structural superposition of dimer AB (chains cA and cB) and dimer CD (chains cC and cD) of EcZraS-CD yields an overall r.m.s. deviation of 2.3 Å ^39^. Chains cA resembles cC while cB has a disposition similar to cD with r.m.s deviation values of 1.0 Å and 0.6 Å respectively (Figure S5A). The r.m.s. deviations between corresponding chains suggest a fairly close structural similarity implying a conserved overall structure, but the r.m.s. deviation value of 1.0 Å suggests presence of minor structural changes between chains cA and cC. Upon close inspection CA domain in cC was found to have rotated through an angle of around 20° (Figure 2C), with a change in the hinge residues spanning Val305-Leu310 in comparison to chain cA ^41^. Besides, structural superposition of cA and cC exhibit a relative change in the base region of the chains, attributed to tilting of the DHp helices (Figure 2A and 2B). We were however unable to trace a short stretch (Ala274-Ala276) of the base loop region in cC of dimer CD.

### Dimers display conformers in an intermediate state resembling a kinase-competent state

The ligand-bound structures of EcZraS-CD display an asymmetric arrangement with the CA domains occupying different planes (Figure 2A). The two CA domains are roughly 36.8 Å and 39.4 Å apart forming angles of around 112.9° and 127.26° respectively between them (Figure 3A and S6A) ^39^. To understand the structural reorientation in the bound states, we modeled an apo form of ZraS (EcZraS-F) comprising of a dimer XY with chains cX and cY (Figure 1C) ^30^. EcZraS-F had a co-planar arrangement of CA domains which are 52 Å apart and form an angle of nearly 180° (Figure 3A). The CA domain of one protomer in EcZraS-CD is located in the vicinity of the DHp bundle, and the other, which is visibly undergoing rotation, is directed away from the core bundle. The average distance between Cα atoms of conserved His254 in the DHp bundle and ATP-binding Asn367 in the CA domain of the opposite protomer are 23 Å and 36.5 Å for the closer and distant domains, respectively, with precise distances being 23.9 Å and 36.7 Å for dimer AB, and 22.2 Å and 36.1 Å for dimer CD (Figure 3D). Helices hα1 and hα1’ of the dimers bend through angles ranging 11-26° at the H-box Pro259 with dimer CD displaying an additional bend N-terminal to the DHp domain towards the linker Lα1 in chain C (Figure 3B). Monomers in EcZraS-F have an equidistant His254-Asn367 Cα distance of around 22.4 Å and a bending angle of nearly 17.5° at Pro259 for hα1 and hα1’ (Figure 3B, 3D). Clustering of available structures of the cytoplasmic domains of SHK based on conserved His-Asn Cα distances, representing relative positioning of CA domains with respect to the DHp core, delineates the role of SHKs based on structural symmetry ^47^. Reportedly, apo and phosphatase-active structures bound to ADP have symmetric arrangements, with both CA domains directed away from the central bundle in an open conformation. Kinase-active structures bound to ATP and its non-hydrolyzable analogs are comparatively asymmetric, with one CA domain associated proximally with the DHp bundle in a closed conformation and the other oriented away in an open conformation.

**Figure 3.**
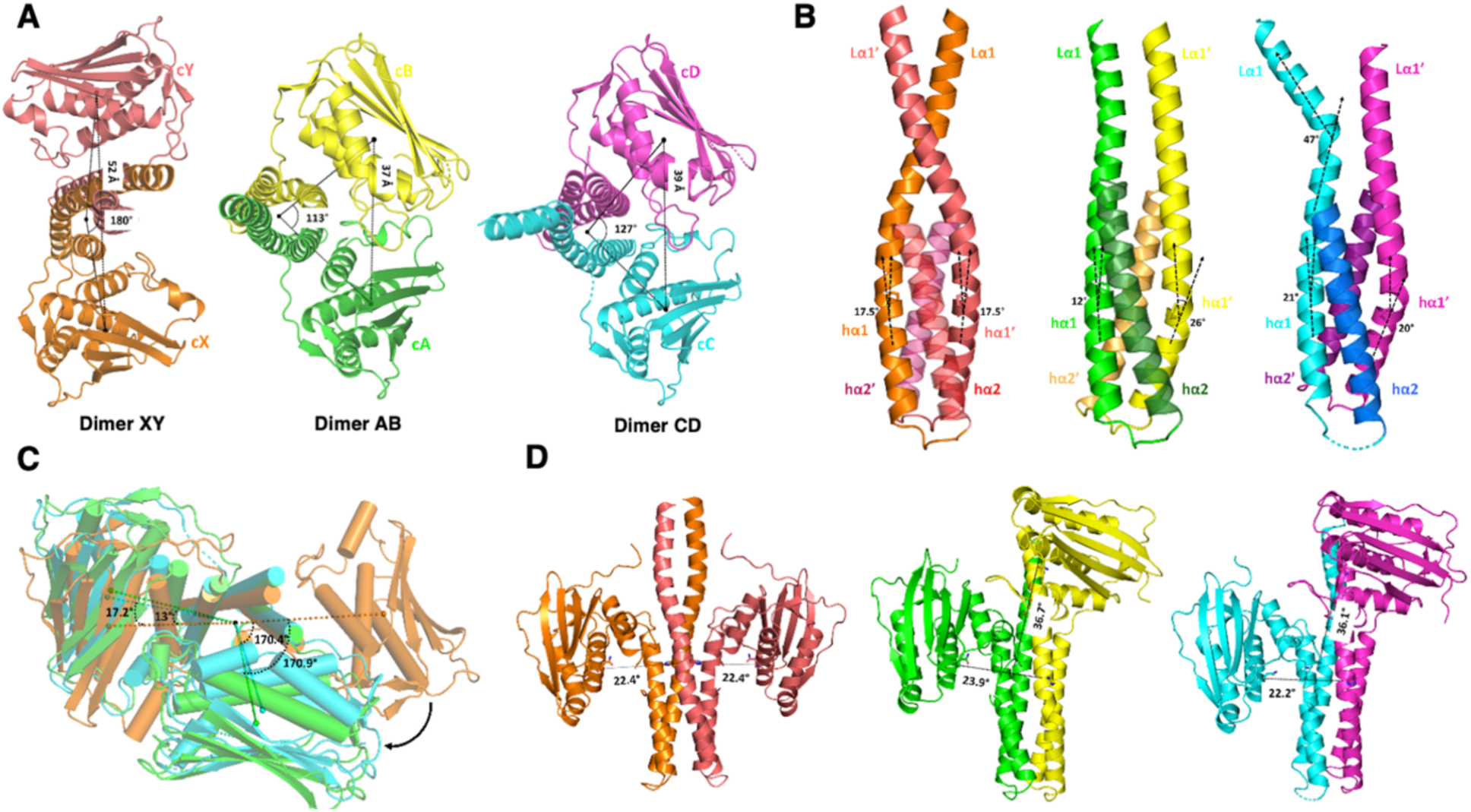
Measured distances and angles of the cytoplasmic domain of ZraS. (A) Inter-domain distances and angles between the CA domains of apo EcZraS-F (dimer XY) and ligand-bound EcZraS-CD (dimer AB and dimer CD) structures. (B) Helix bending angles at Pro259 of helix hα1 (orange) and hα1’ (deep salmon) of dimer XY, helix hα1 (green) and hα1’ (yellow) of dimer AB, and helix hα1 (cyan) and hα1’ (light magenta) of dimer CD. (C) Structural superposition of dimer XY (orange), dimer AB (green), and dimer CD (cyan) displaying angular rotation of CA domain upon ligand binding. (D) Distance between catalytic His254 in the DHp and ATP-binding Asn367 in the opposite CA of apo (dimer XY) and ligand-bound (dimer AB and dimer CD) structures.

Michaelis-complex forming CpxA (4CB0) and phosphorylated DesK (5IUM) exhibit CA domains in varied planes with a single CA domain in closed conformation owing to steric hindrance ^10,19^. Formation of closed-state CA-DHp arrangements is simultaneously coordinated with helical bending in the DHp bundle of WalK and CovS through 25.7° at Pro396 and 25° at Pro222, respectively ^13,14^. Intermediate states possessing similar coordinated bending at the DHp helices in both protomers through small angles were speculated for WalK during autophosphorylation ^14^. The distance between each proximally positioned CA domain and the DHp bundle in the ADP-bound dimers of EcZraS-CD ranges 20–30 Å. It is comparable to known ADP-bound structures of SHKs but insufficiently large, considering a distance of 10–15 Å required for a closed-state autophosphorylation complex formation ^47^. The remarkable structural asymmetry observed for the obtained dimers with considerable helical bending and relative positioning of CA domains, suggest the presence of dimers in two different intermediate conformations approaching a kinase-active state.

### Nucleotide-binding pocket

A nucleotide-binding pocket in EcZraS-CD is formed between helices α2 and α3, delimited by sheets β6 and β7. It is enclosed by a loop forming the ATP lid. ADP and Mg^2+^ lie in the pocket lined by N, G1, F, G2, and G3 box residues of the CA domain (Figure 4A). Amino acid residues from the G1-box (green cyan) form the base of the pocket. The roof region is lined by residues from G2-box (forest), Gripper region (marine), and N-box (deep teal). The G2-box flanks the pocket bordering one side occupied by the two phosphate groups of ADP. The opposing side is enclosed by residues from the N-box. The back wall is formed by residues of the G1-box (green cyan), a patch of G-3 box (gray40), and F-box (cyan). ATP lid loop, a region distal to the F-box and proximal to the G2-box, forms the access door to the pocket, extending towards the pocket base. The N6 amino group of the adenine ring in ADP forms a hydrogen bond with Asp395 of G1-box. An additional interaction responsible for nitrogenous base binding includes a long-range cation-π interaction between positively charged Lys416 and the adenine ring ranging 5.5-6.5 Å. Asp395, Ser448, Gly399, and Gly397 form the groove accommodating the adenine ring of ADP. 2’ and 3’ hydroxyl oxygen atoms of ribose sugar interact with side chain oxygen atoms of hydroxyl groups and main chain nitrogen atoms of ATP lid loop residues Thr414, Thr415, and Lys416. The α and β phosphate group oxygen atoms are bonded to Thr420 of the loop region and Leu424 of the G2-box, which assists in the orienting ADP inside the pocket. The residues Phe409 and Phe413 of F-box do not interact directly with the nucleotide but flank a side near the ATP-lid loop region. Oxygen atoms of the conserved residue Asn367 of N-box and phosphate groups of ADP coordinate with Mg^2+^ (Figure 4B). SHK structures obtained in high resolution have an octahedral Mg^2+^ binding geometry with closely coordinated water molecules. Due to the low resolution of the obtained structure, water molecules could not be modeled despite the presence of positive electron density in the pockets after fitting the nucleotide and Mg^2+^.

**Figure 4.**
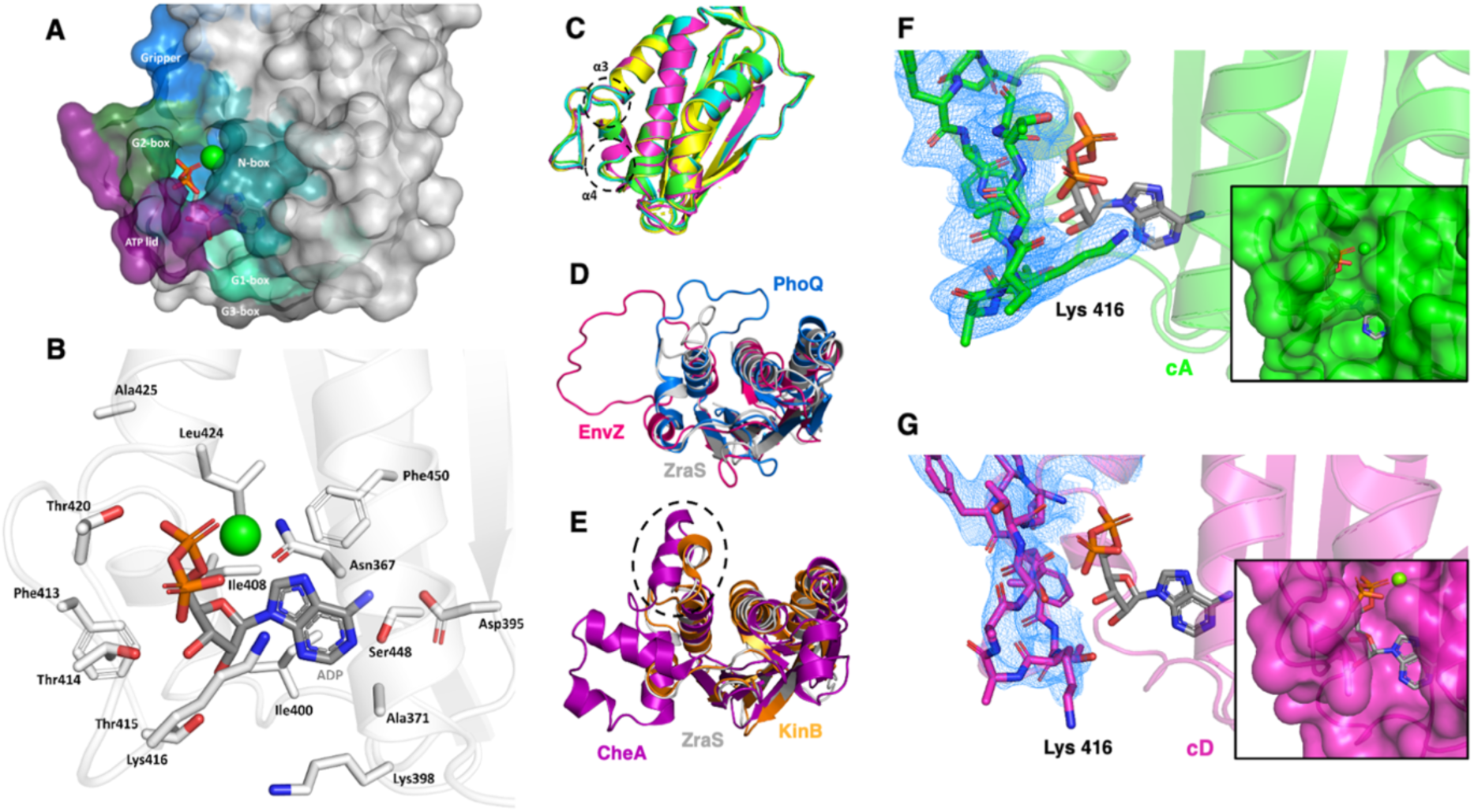
Nucleotide binding pocket and lid loop of EcZraS-CD. (A) Surface representation of the nucleotide-binding pocket lined by residues from N (deep teal), G1 (green cyan), F (cyan), G2 (forest), and G3 (gray50) boxes in EcZras-CD. (B) ADP and amino acid residues flanking the nucleotide-binding pocket of cA are shown in stick representation. Mg^2+^ is shown as a sphere (green). (C) Structural superposition of CA domains of chain A (green), chain B (yellow), chain C (cyan) and chain D (light magenta) of ZraS showing local uncoiling of α3 and α4. (D) Structural superposition of ATP lid loop region of PhoQ (marine) in a closed-state and EnvZ (hotpink) in an open-state with EcZraS-CD (gray90). (E) Comparison of ATP lid loop region of CheA (purple) and KinB (orange) with EcZraS-CD showing secondary structures in the loop. (F) and (G) Stick and surface representation showing change in Lys416 interaction in cA and cD. The dynamic lid loop encloses the pocket in cA interacting with adenine ring of ADP, absent in cD. The 2FoFc map is contoured at 1.0 σ.

### A flexible ATP lid loop guards the nucleotide binding pocket in EcZraS-CD

The ATP lid loop of EcZraS-CD is a 14-residue stretch consisting of residues Thr410-Gly423, bordered by helices α3 of G2-box and α4 of F-box. It is anchored at one end by the conserved Phe409 residue of helix α4 and the other by a conserved hydrophobic residue Leu424 of helix α3. Phe409 is inserted into a characteristic hydrophobic patch of SHKs, formed by residues Ile408, Leu424, and Ile438. The loop region is neither directed towards the core similar to the long ATP lid loop of PhoQ (1ID0) nor directed away as in EnvZ (1BXD) (Figure 4D) ^11,15^. The pocket and lid loop form an ordered structure as opposed to locally disordered nucleotide-free structures of NR(II) (1R62) and CpxA (4BIX) ^10,48^. The region’s mobility is regulated by conserved glycine residues of the G2 box (Gly419, Gly421, and Gly423). Structural comparison with CheA and PhoQ having loop lengths of 19 and 16 residues, respectively, shows a relatively shorter loop region for EcZraS-CD ^15,46^. Being relatively short it stays in an intermediate position and encloses the nucleotide binding pocket upon interaction of the lid loop with the DHp helices. Local unwinding in helices α3 and α4 help regulate the loop length upon nucleotide binding (Figure 4C). Though no drastic change in the conformation of the loop region was observed, a change in interaction of Lys416 with the ADP molecule enclosing the pocket in a proximally located CA domain of cA was found, absent in the distally located CA domain as in cD (Figure 4F and 4G). Change in lid loop conformation favoring nucleotide binding upon interaction with DHp bundle residues was observed for sensor kinases ^49^. Other than acquiring different conformations upon nucleotide binding, the ATP lid loops display presence of secondary structures in CheA (1I58) and KinB (3D36) (Figure 4E), absent in the lid loop region of EcZraS-CD ^46,50^. The CA domain of EcZraS-CD resembles the CA domain of CckA bearing 35% sequence identity (Figure S4A and S4B), a highly conserved ligand-binding pocket and the signature TTK lid loop motif but lacking the cyclic di-GMP binding residues of CckA ^51^.

### DHp helices undergo a rotary switch to transition between alternate conformations

The ability to accommodate different types of signaling needs is reflected through the diversity of DHp bundles found in SHKs. Dimerization specificity and alignment being indispensable for their catalytic function, disruption in the bundle through mutations and deletions, prevent kinases from effectively performing autophosphorylation, phosphate transfer and phosphatase activities ^52,53^. Depending on sequence conservation and its location, the DHp bundles have been classified into HisKA (PF00512), HisKA_2 (PF07568), HisKA_3 (PF07730), and HWE (PF07536) domains in the Pfam database. EcZraS with an H-box consensus of His-Glu-Ile-Arg-Asn-Pro-Leu falls under the category of HisKA bundles. HisKA bundles are commonly made of two helix-turn-helix motifs, arranged in an antiparallel manner with both turns at one end, while the N and C-termini are exposed towards the other end.

Analysis of the bundle in apo EcZraS-F using programs such as TWISTER ^42^ and SamCC Turbo (manual mode) ^43^ helped identify the distribution of residues (heptads) and their layering pattern, roughly spanning Ala253-Phe270 and Leu301-Ala284 in both chains of the dimer. The chains assemble into a heterotetrameric A2B2 coiled-coil possessing a hydrophobic core made up of “a” and “d” residues (Figure 5A). These residues protrude out along the helices facing their counterparts in the diagonally opposed helix arranged in multiple alternating ad layers, by forming knobs housed inside cavities or holes produced between knobs of nearby helices. Position “d” is occupied by large hydrophobic residues such as Leu, Ile, and Met and “a” by small residues like Ala with an occasional presence of Leu and Val. Helix hα2 and hα2’ are translated downwards compared to hα1 and hα1’ by roughly half a turn, resulting in a staggered arrangement of layers as opposed to parallel layers found in conventional coils. The offset bundle packing has an alternating disposition of knobs constituting the inner bundle core predominantly occupied by “d” residues of the ad layers (adL1-adL6), having Ile in helices hα1 and hα1’ and Leu in helices hα2 and hα2’ (Figure 5C). “a” residues are found lining the inner core made by side chains of “d” residues (Figure 5B and 5D).

**Figure 5.**
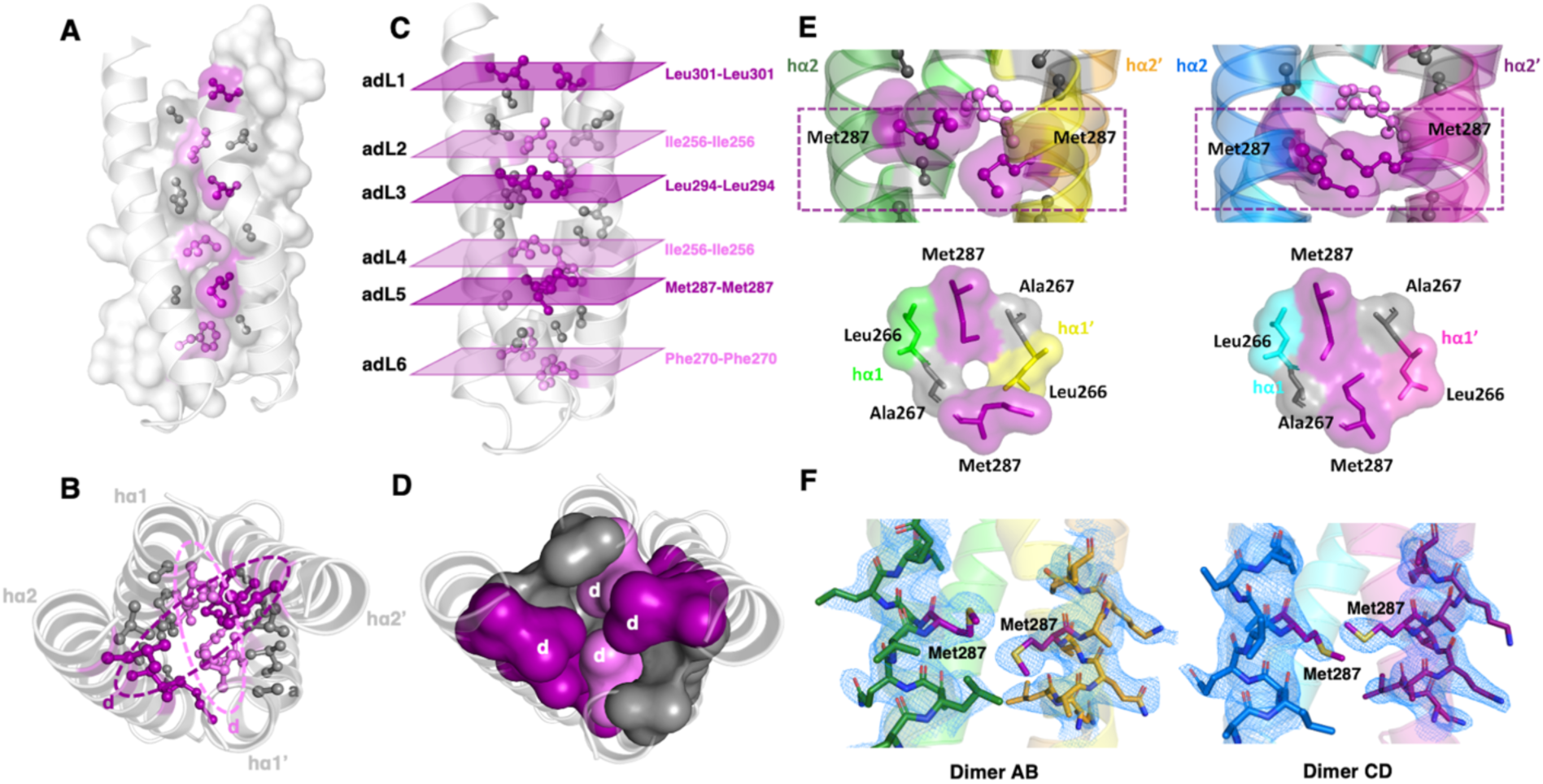
Organization and core packing of DHp bundle in EcZraS-CD. (A) Representation of DHp bundle forming modified knobs into holes packing. Knobs formed by sidechains of residues at position “a” (gray50) and “d” (purple and violet) are packed into grooves formed between ridges of neighboring helices along the dimer interface. (B) Alternating residue layers (adL1-adL6) at position “d” from helix-1 (purple) and helix-2 (violet) forming the inner core of DHp bundle. (C) The top view of the bundle showing the distribution of “a” and “d” residues forming the a and d layers. (D) Surface representation of “d” residues (violet and purple) in the inner core and “a” residues (gray50) lining its periphery. (E) Lateral and top view of variable interactions between Met287 in adL5 at the dimerization core interface of dimer AB and dimer CD. (F) 2FoFc electron density map contoured at 1.0 σ showing Met287 in the DHp bundle of dimers AB and CD.

TWISTER identified a similar heptad distribution for the bundles in ligand-bound EcZraS-CD dimers. Owing to local changes induced upon ligand binding, SamCC Turbo was unable to detect the corresponding residue stretches automatically. In order to determine the layering pattern and helical parameters, we defined the bundle stretches manually using heptad positions defined by TWISTER. The layering pattern in EcZraS-CD dimers remained consistent, having visible fluctuations in parameters such as Crick angle deviation. With a rather similar conformational disposition, the DHp bundle layers were found to have different Crick angle deviation values per layer in dimers AB and CD, signifying a change in their core packing geometry supported by helical rotation (Table S1-S3 and Figure S8-S10). Other helical parameters coupled to helical rotation in four-helix bundles, such as axial shift and axial radius have also been measured (Table S4-S7 and Figure S11-S14).

Comparison of r.m.s.d values of regions across the DHp bundle show minimal deviation in the mid region (Figure S5B and S5C). Scanning for repeats in the linker regions of dimers AB and CD, the program TWISTER helped identify a region containing an insertion (stutter) juxtaposed between the linker (L*α*1/ L*α*1’) and DHp (h*α*1/ h*α*1’) helices (Table S8). Amino acid insertions play an important role facilitating conformational transition in numerous kinases ^10,12^. Structural changes caused as a result of deviations is accommodated across the length of the helical continuum (accommodation length) ^54^, resulting in bundle repacking during signal transmission from the transmembrane to the kinase domain. Presence of stutters leads to an occurrence of non-heptads amongst heptads allowing divergence from the traditional knobs-into-holes packing of the core forming positions “a” and “d”. The “d” residues in dimers AB and CD point straight toward the central axis forming tip-to-tip interactions, or are directed sideways, enclosing the central core. Figure 5E and 5F shows one such “d” residue (Met287) packing of in the layer adL5. The sideway interaction of Met287 with Ala267 in dimer AB is replaced by nearly a tip-to-tip interaction between Met287 of each protomer in dimer CD. Rearrangement of residue packing between different helices at the dimerization core interface of dimer AB and CD, leading to the reorganization of knob placement into the holes, reveals the implication of core packing during conformational alteration. Residue interactions at different helical interfaces of the DHp bundle in EcZraS-CD dimers have been shown in Figure S7A and S7B.

### Variable interactions between polar and hydrophobic residues in the DHp bundle interface lead to changes in the helical registry

The four-helix bundle of EcZraS-CD is stabilized through hydrophobic interactions of the knob forming “d” residues at the inner core of the DHp bundle, guarded by “c” and “g” residues forming helical interfaces at the periphery between hα1-hα2’ and hα1’-hα2. Upon intervention and differential interaction of locally interspersed predominantly polar residues such as Glu255, Ser262 and Glu290 at positions “c” and “g”, dimers AB and CD exhibit alterations in their core layering arrangement (Figure S8A and S8B). The presence of polar residues evades formation of a tightly packed core, a requisite for underlying bundle flexibility in proteins. Dimer AB and CD having two variable conformational protomers exhibiting unlike interactions at the peripheral interface between helices hα1-hα2’ and hα1’-hα2. Interface of hα1’-hα2 lying closer to the proximally located CA domain exhibit prevalence of c-c and g-g interactions in contrast to the c-g pattern observed at the distant interface of hα1-hα2’, demonstrating a change of helical registry as in Dimer AB (Figure S8C). Gomez et al. emphasized on the importance of core and peripheral residue interactions in controlling the stability and plasticity of coiled-coils ^55^. Translational motion in the coiled coil stalk bundle of the motor protein dynein is regulated by balancing the strength between hydrophobic and electrostatic interactions at the core and peripheral interfaces of the superhelix ^56^.

### Aromatic amino acid interactions and loop residues at the DHp base regulates the directionality of bundle rotation

Autophosphorylation of SHKs in cis (left-handed) or trans (right-handed) mode requires modulation of the CA domain motion across the DHp bundle, aided by changes in the bundle base region ^18,57^. In vitro biochemical studies in *E. coli* identified trans-autophosphorylation in ZraS, with inadequate evidence on the structural basis of its regulation ^58^. The analysis of asymmetric dimers AB and CD of EcZraS-CD shows distant CA domains undergoing vertical shift combined with a rotation of nearly 170°, while the proximal CA domains rotated with very minimal variations (Figure 3C). The hydrophobic inner core of the dimeric DHp bundles with predominantly aliphatic residues, have a localized distribution of aromatic amino acids such as Tyr269 and Phe270, towards the bundle base region of helices hα1 and hα1’. Sidechains of Phe270 residues are projected towards the inner core and flanked by solvent-exposed Tyr269 on either side of the homodimer. In dimer AB, Phe270 residues from cA hα1 and cB hα1’ are placed 4.8 Å distant, oriented in a parallelly displaced manner.

Comparison of spatial positions of Tyr269 and Phe270 indicate presence of Tyr269 close to Phe270 in cA separated by a distance of 5.0 Å, while being positioned 5.7 Å apart in cB (Figure 6A). Tyr269 residues further exhibit varied interactions with the bundle base residues and base loop bordering Glu279 residues, resulting in two different interfaces between base regions of hα1-hα2’ and hα1’-hα2, with only Tyr269 in cA interacting with Glu279 in cB. A comparable conformation resembling that of dimer AB is also observed in dimer CD confirming an adaptable aromatic π-π stacking in Tyr269-Phe270 and electrostatic interactions in Tyr269-Glu279 on either sides of the dimeric bundle, potentially generating a torque aiding bundle rotation (Figure 6B). Furthermore, an untraceable loop region in cB confirms the dynamic nature of the bundle base.

**Figure 6.**
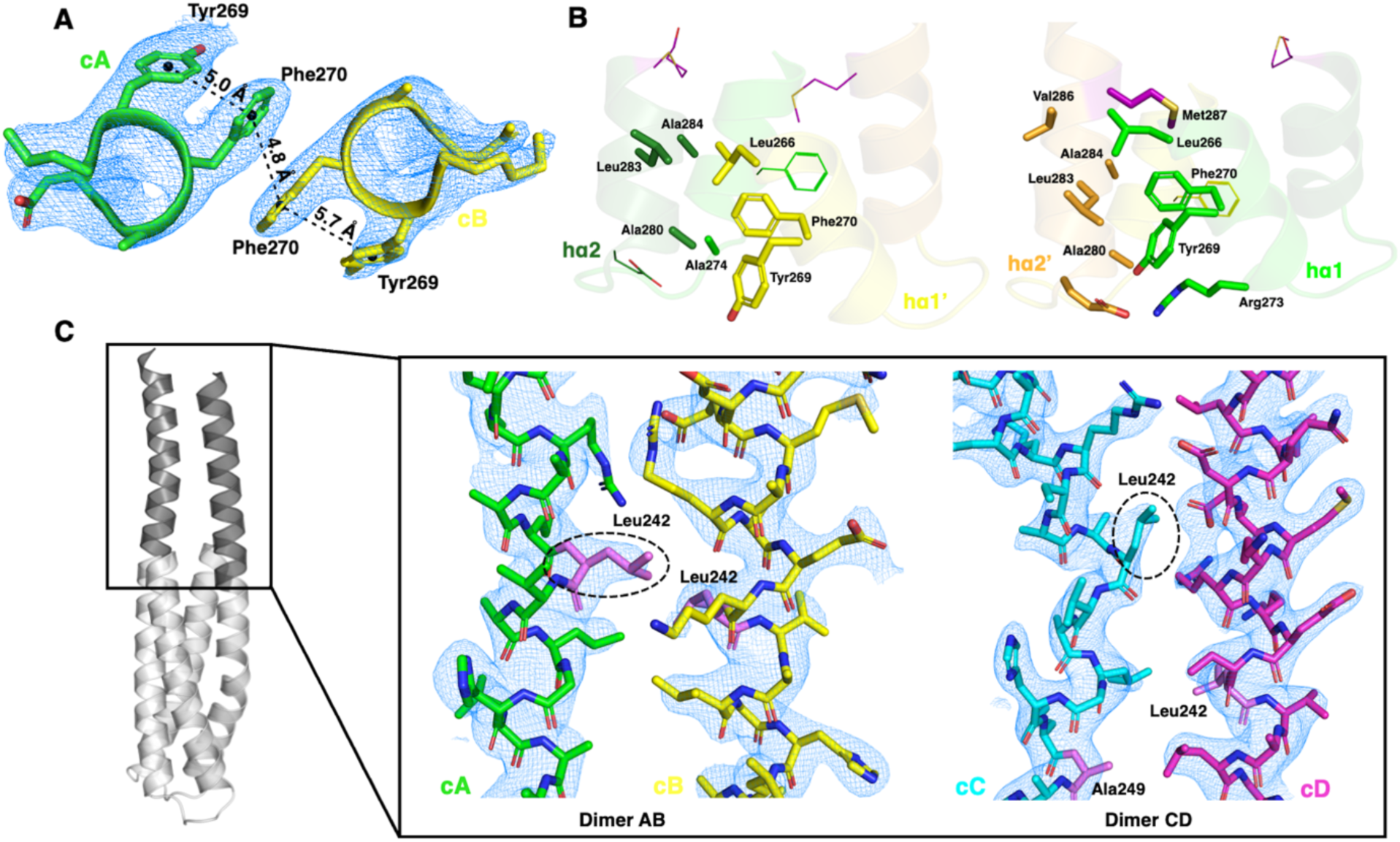
Variable helical interface interactions of DHp and linker domain in EcZraS-CD. (A) Distance between aromatic amino acid residues Tyr269 and Phe270 of chains cA and cB in dimer AB. (A) Bundle base interactions in dimer AB regulating bundle rotation. Interacting residues are shown in stick representation and non-interacting residues in line representation. (C) Residue interactions at the linker interface of dimer AB and CD. Distortion in the alternate layering arrangement of the core residues is caused due to intervention by residues in polar clusters. The core “d” residues interacting at the interface in each dimer are labelled in violet with the 2FoFc maps showing their occupancies, contoured at 1.0 σ. The dotted circles highlight the position of linker helix bending initiating at Leu242 in cA and cC.

### Conserved helix disruptors facilitate the scissoring of helices in the core bundle

Residues such as Pro and Gly are known disruptors of α-helices ^59–61^. They are commonly found in helical regions requiring flexibility and undergoing mobility. Pro259 in the H-box of the DHp bundle forms a kink in the backbone helix of EcZraS-CD. Regions N and C-terminus to Pro259 in the dimers AB and CD bend variably with minimal deviation in the stretch containing the residue. The helices hα1 and hα1’ in dimer AB bends through angles of nearly 12° and 26° and that of dimer CD through 21° and 20° (Figure 3B). Helices present in both membrane and water-soluble environments, as in signaling helices of GPCRs and SHKs, stalk regions of motor proteins, and helices involved in ion-channel gating rely on kinks or hinges to carry out their biological functions ^13,62–64^. The majority of SHKs possess a conserved Pro kink, with the exception of a Gly kink found in some, such as DesK (3GIG) ^12^. A study on VicK established the importance of Pro mutants in controlling phosphatase activity ^13^. Point mutation from Pro to Ala eliminated phosphatase activity that could be rescued by replacing it with a Gly, suggesting the necessity of a hinge-forming residue in state transition.

### A unique linker relays signal from transmembrane helix TM2 to hα1 and hα1’ of DHp bundle

The cytoplasmic helical region contiguous with the kinase domain extends N-terminally to form the linker domain, connecting the membrane-passing helix TM2. The linker region in EcZraS-CD lacks other intermediate signal transducing elements featuring tertiary folds and rather comprises of a simple helical structure differing from well-studied α-helical SHK linkers such as S and Jα-helices ^65^. Sequence analysis using psi-BLAST, pLM-BLAST, PLM Search and multiple sequence alignment with well-studied S-helices failed to identify any structurally characterized SHK homologue. Presence of a coiled-coil architecture could not be confirmed using SamCC Turbo. Conversely, TWISTER identified presence of both canonical and non-canonical coiled-coil repeats interspersed by a stutter. Residues at position “d” of the region are primarily occupied by hydrophobic residues, with “a” occasionally occupied by polar residues. In absence of a precise boundary delineation, the α-helical linker and the DHp helices hα1 and hα1’ coiled-coils closely overlap along their length. The linker stretch consists of polar residue clusters Arg227-Gln230 (RSRQ) and Gln233-Met236 (QDEM) that reduce the energetic stability of the coiled-coil interface, favoring structural plasticity. Comparison of linker interfaces of dimer AB and CD shows differences in bundle interactions during signal transduction (Figure 6C). cC adopts a more curved configuration bending through an angle of nearly 47°, attributed to the interactions with polar residues such as Arg238 and Asp234.

### CA domain in the vicinity of the DHp bundle forms a CA-DHp interface using ATP lid loop residues

CA domains of cA and cC in EcZraS-CD form CA-DHp interfaces with hα1 of cB and cD, respectively. ATP-lid loop residue Tyr412 of the proximal CA domain, is seen to primarily interact with the DHp residues of the opposite protomer in the dimers AB and CD. It interacts with DHp residues Leu248, Gly251 and Val252 (Figure 7A). Pro411 of cA and cC interacts with Tyr412 and Leu248 residues of the CA domains of cB and cD. No visible interactions were observed between CA residues and His254 alike previously known Michaelis states ^10,18^. Electrostatic surface potential analysis shows the interface interactions are an outcome of electrostatic complementarity between a positively charged surface in the CA domain and a localized negatively charged surface at the DHp bundle (Figure 7B). Additional CA-CA residue interactions were visualized for dimer AB, absent in dimer CD. These interactions could be essential for the functional coupling of the domains, where changes in the conformation one CA domain affects the conformation and activity of the other thereby aiding allosteric communication during the process of autophosphorylation. The relative positioning of the CA domains during autophosphorylation could potentially be regulated in a single-step process through direct interactions with the complementary DHp bundle residues or through a multi-step process involving the formation of several transitional interfaces using diverse sets of residues in either domain. The absence of direct contact between the CA domain residues and DHp His254 in the dimers of EcZraS-CD, suggests the process involves a multi-step transition with gradual changes through internal rearrangements and domain-domain allostery.

**Figure 7.**
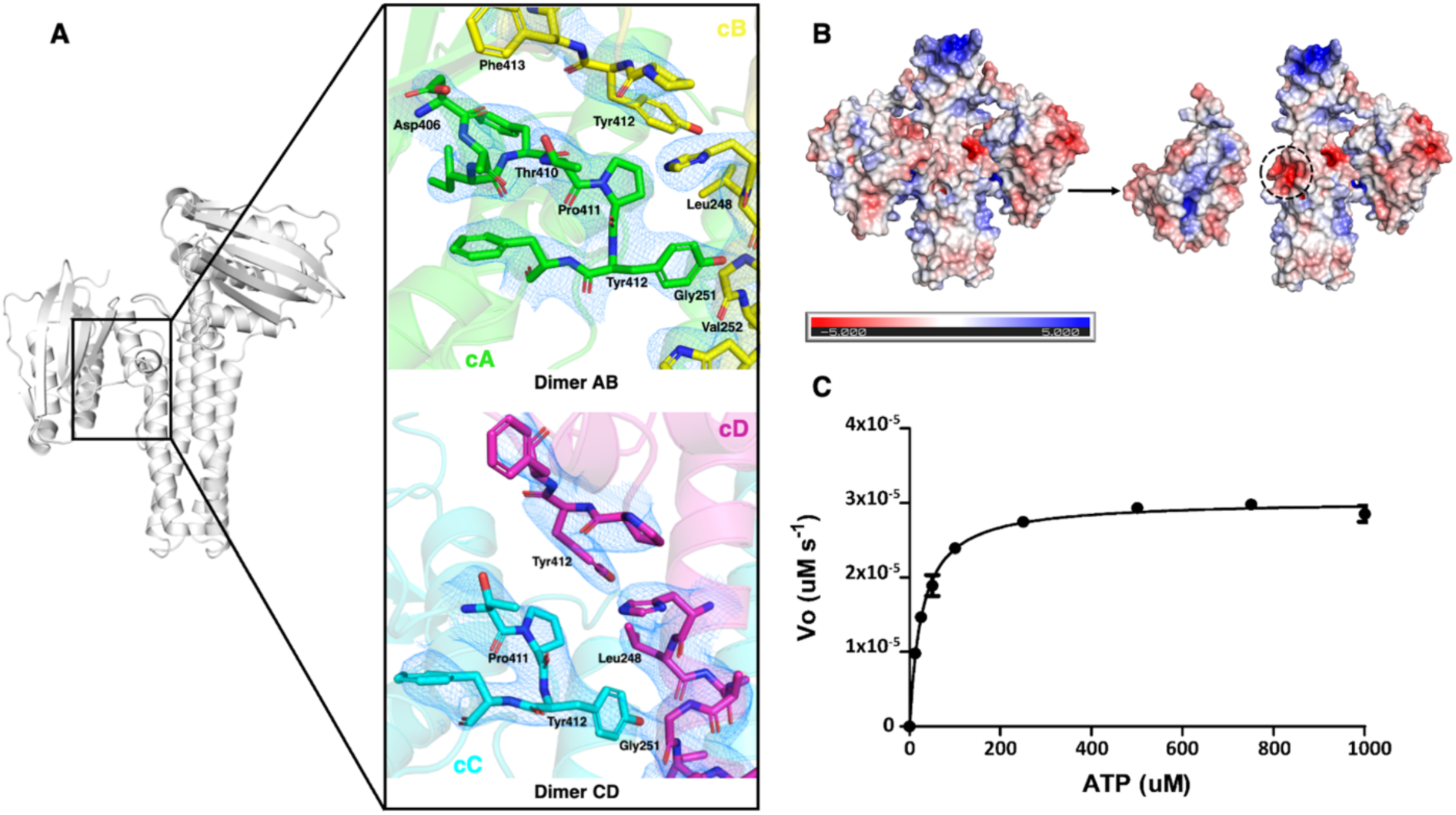
Proximal CA-DHp domain interfaces and enzyme kinetics plot of EcZraS-CD. (A) Cartoon representation of CA-DHp interface of dimer AD and dimer BC. Sticks show the interacting residues at the interface in either dimer. (B) Electrostatic surface potential representation of interacting interfaces of CA and DHp domain. ADP bound to the nucleotide-binding pocket, with a highly positive electrostatic potential is shown in surface and stick representation. (C) Kinetic characterization of EcZraS-CD autophosphorylation using Pyruvate kinase-lactate dehydrogenase coupled assay. The curve shows ATP dependence on initial velocities of autophosphorylation.

### Kinetic analysis of EcZraS-CD

The rate of autophosphorylation of EcZraS was estimated using coupled enzyme kinetics (Figure 7C). The K_m_ and V_max_ values were determined to be 27.31 and 3.032 x 10^-5^ uM/s with an enzyme concentration of 15uM.

## Conclusion

Initiation of signal transduction through autophosphorylation is a cumulative outcome of input enhancement and reduction effects by multiple domains in SHKs. The constituent domains undertake numerous changes leading to transmission of signaling information across the bacterial membrane. To study the alterations central to conformational transition during these signaling events, we determined the structure of the truncated cytoplasmic stretch of ZraS from *Escherichia coli*, bound to its ligand (ADP) and Mg^2+^.

The overall structure of EcZraS-CD consists of a functional homodimer with the conventional kinase domain, continuing N-terminally to a unique α-helical linker domain. Two dimers were obtained in the unit cell of the crystal, trapped in successive conformational states undergoing state switch. Their asymmetric arrangement is in accordance to the trans mode of autophosphorylation, observed in *in-vitro* biochemical studies ^58^. A comparison made between the two conformers provide evidence on the reorganization of the malleable bundle interface through helical rotation, helical bending, shift in helical registry and variable accommodation of amino acid insertions along the length of the helices during conformational alteration. The dimeric kinase bundles present a reorganization of the hydrophobic inner core layers due to intervention of polar residue interactions favoring rotary switch. The rotational switch in conformation is simultaneously aided by amino acid insertions in the helical linker-DHp backbone variably accommodated across the backbone length. Flexibility of the bundle is further enhanced by the presence of a hinge forming Pro259, allowing helical movements analogous to scissors (scissoring) and facilitating twisting of localized helical regions in different orientations about the hinge (swiveling).

The linker region forms a coiled-coil dimer that deviates from known signaling helices, exhibiting pronounced helical bending and a change in interface interactions. The dimers also offered insights into how the directionality of rotation is regulated, linking it to the change in base and loop residue interactions. Based on the peripheral helix interfaces and spatial orientation of the residues, it is plausible that the torque required for rotation generated at the bundle base through differential interactions between aromatic residues (Tyr269 and Phe270) and the loop bordering residue Glu279. The notable bending at the base region of cC with a base loop whose conformation could not be traced due to its dynamic nature as compared to its corresponding chain cA, suggests that changes in loop conformation have a role in regulating the directionality of bundle rotation. Evaluation of the residue interactions at the CA-DHp and CA-CA interfaces shows positioning of the proximal CA domain near the H-box to be a multi-step process, guided by the DHp helix and the opposite CA domain, thus validating the concept of inter-domain allostery. The CA-DHp interface of EcZraS-CD lacking any direct interaction with the phosphorylating His254 residue suggests the dimers could be potential occluded state approaching a configuration critical for autophosphorylation.

Information transfer in sensor kinases such as ZraS is indeed a much more elaborate and intricate process, with sequential and topological diversity furthering its complexity. A multitude of well-orchestrated changes helps them surf across conformational landscapes in a sequential manner, traversing a spectrum of numerous conformers (Figure 8). Future studies capturing additional conformations will be necessary to determine the precise sequence of events leading to signaling. Nonetheless, our results present substantial evidence on the occurrence of intermediate states and combination of structural changes carried out during signal transduction in ZraS.

**Figure 8.**
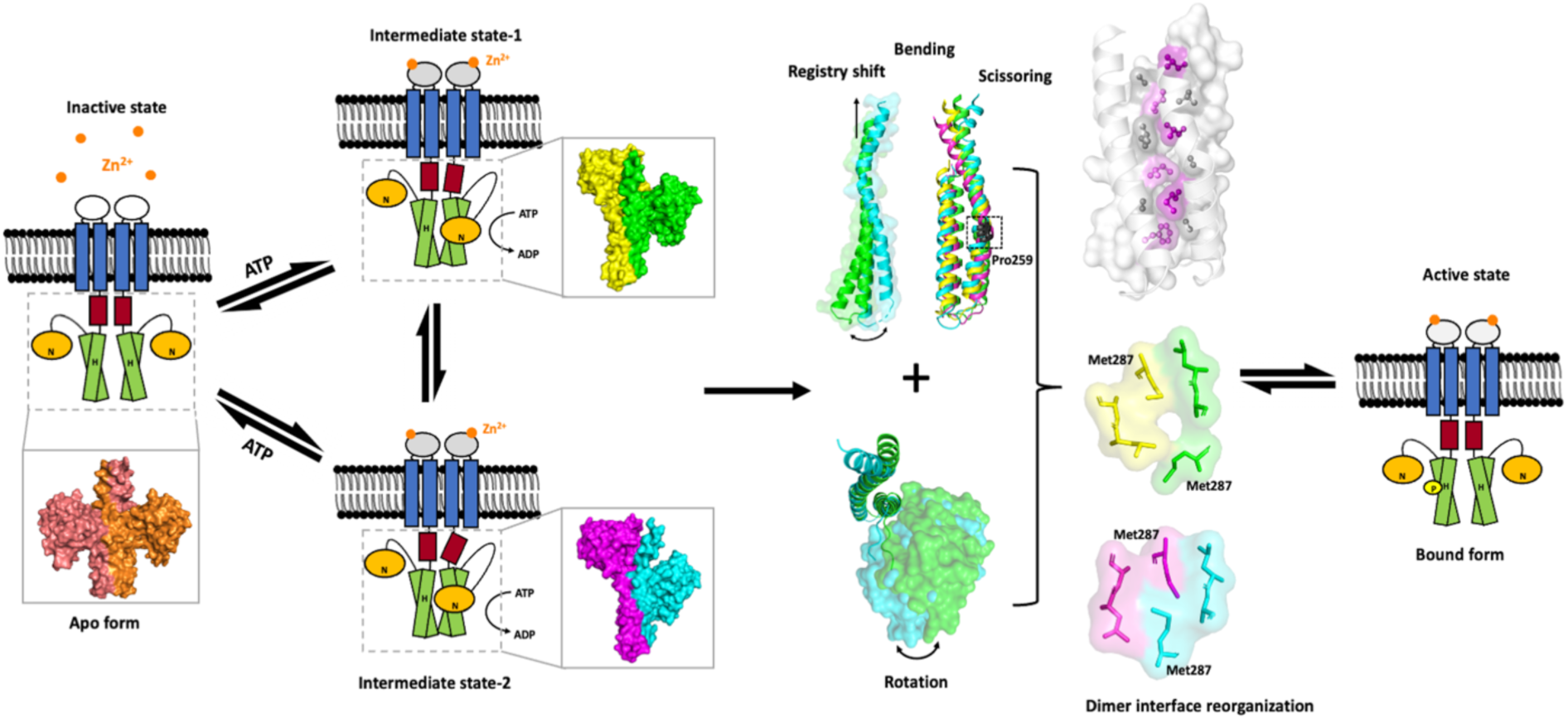
Model of autokinase activation in EcZraS-CD. Schematic representation of the autokinase activity in the cytoplasmic domain of ZraS illustrates transition from an inactive apo form to an active ligand-bound form mediated through intermediate conformers, state-1 (dimer AB) and state-2 (dimer CD). Conformational transition in the cytoplasmic region is facilitated through dimer interface reorganization driven by helical bending, scissoring, registry shift and domain rotation.

## Author contribution

R.A. conceptualized and designed the research; N.M., P.M., and R.A. executed the research. N.M. prepared the samples and performed biochemistry. P.M. grew the crystals. R.A. collected the X-ray diffraction data, processed the data, and solved the structure. N.M. refined the structure. S.P. helped in the initial refinement. N.M. wrote the original manuscript draft. R.A., writing-review and editing the manuscript. N.M. and R.A., visualization. R.A., supervision; R.A., project administration.

## Acknowledgment

We thank Prof. William F. DeGrado for providing the clones of full-length *E.coli* ZraS, reading the manuscript and suggestions.

## Funding and additional information

N.M., P.M., S.P., and R.A. thank NISER for funding and the NISER X-ray diffraction facility. The X-ray diffraction datasets for EcZraS-CD crystals were collected at beamline ID30A ESRF Grenoble. The data collection at ESRF was supported by the grant DBT BT/INF/22/SP22660/2017. N.M., thank CSIR for fellowship. P.M., thank DBT for the DBT-RA fellowship.

## Abbreviations

ESR: Envelope stress response
TCS: Two-component system
SHK: Sensor histidine kinase
HPK: Histidine protein kinase
RR: Response regulator
DHp: Dimerization and histidine phosphoacceptor domain
Hpt: Histidine-containing phosphotransfer domain
CA: Catalytic domain
ATP: Adenosine triphosphate
ADP: Adenosine diphosphate
BME: Betamercaptoethanol
Pfam: Protein family database
DNA: Deoxyribonucleic acid
PMSF: Phenylmethylsulfonyl fluoride
TEV: Tobacco Etch Virus
CC: Coiled-coil
XDS: X-Ray detector software
NADH: Nicotinamide adenine dinucleotide hydrogen
NAD+: Nicotinamide adenine dinucleotide
PK: Pyruvate kinase
LDH: Lactate dehydrogenase
PDB: Protein data bank
APBS: Adaptive Poisson-Boltzmann Solver.

## Ethics statement

The authors have nothing to report.

## Data availability

All the data generated or analyzed during the study have been included in this article and its supporting information. The coordinates of crystal structure deposition have been in protein data bank with PDB ID **9JFJ** (Table 1).

## Supporting information

Additional supporting information can be found online in the Supporting Information section.

