## Supplemental Figures S1 - S14 and Table S1-S6 for "Insights into the conformational dynamics of the cytoplasmic domain of metal-sensing sensor histidine kinase ZraS"


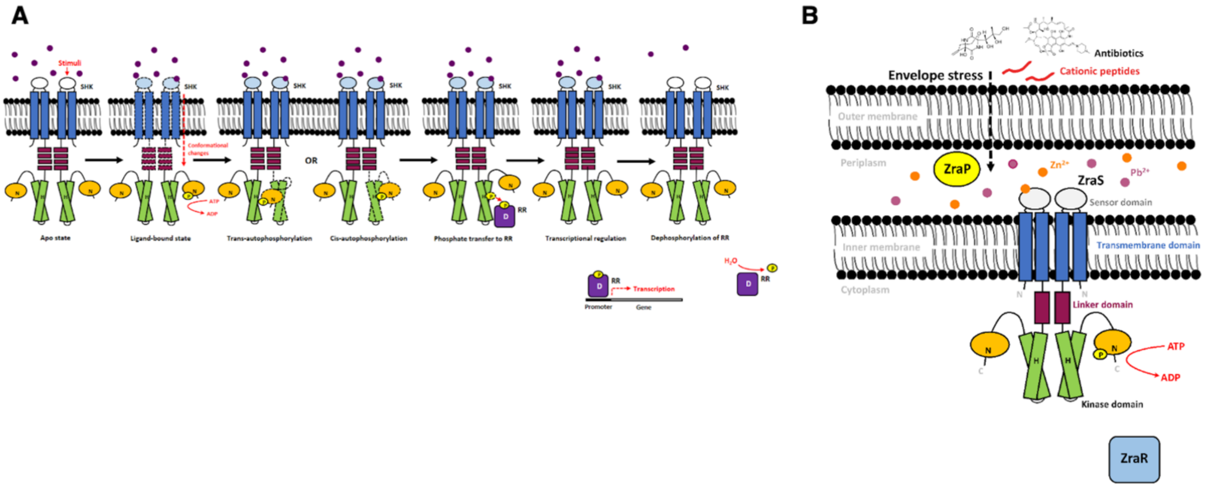


**Figure S1. Stimuli, domain organization and mechanism of signal transduction in SHKs:** (A) Schematic representation depicting the flow of signal transduction in SHKs, including autophosphorylation of SHK (cis and trans modes), phosphate transfer to RR, and dephosphorylation of RR. (B) TCS encoded by operon zraPSR in *Escherichia coli*. ZraS is a membrane-bound SHK that senses envelop stress, coupled with its cognate response regulator, ZraR. ZraP is a periplasmic protein known to function as a zinc chaperon.

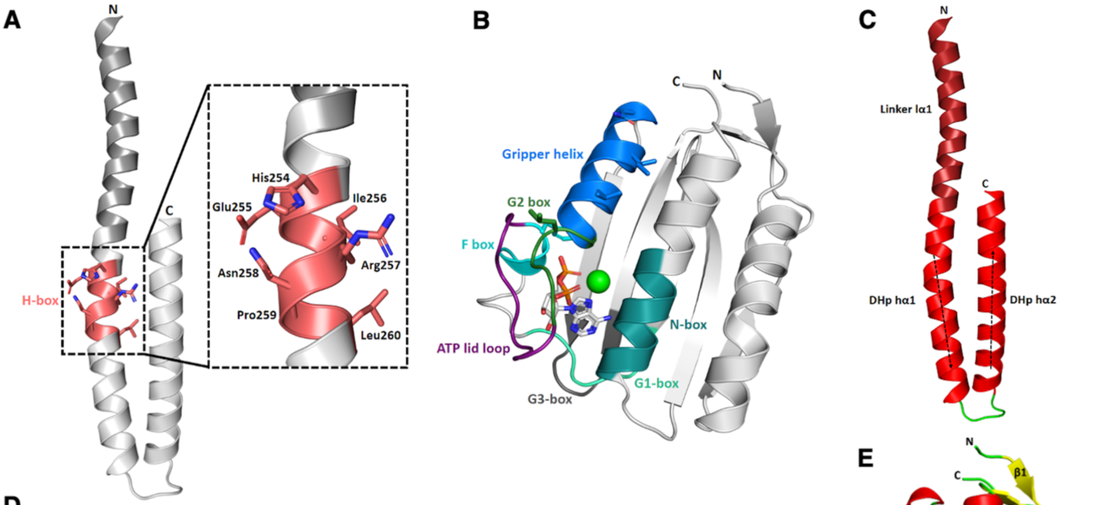



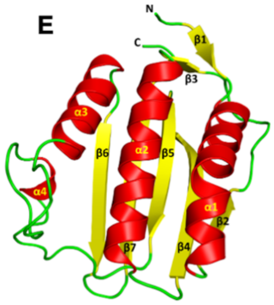



D

| Box | Sequence | Conserved residues | Location |
|---|---|---|---|
| H (His) | HEIRNPL | His254 | hα1 of DHp |
| N (Asn) | NLYLNAIQA | Asn367 | α2 of CA |
| D (Asp) and G1 (Gly) | VTDSGKGI | Asp395, Gly397, Gly399 | βE and adjacent loop of CA |
| F-box (Phe) | DAIF | Phe409 | α4 of CA |
| G2-box (Gly) | GTGLG | Gly419, Gly421, Gly423 | Loop adjacent to α3 of CA |
| G3-box (Gly) | SQEGKGS | Gly447 | βF, βG and βF-βG loop of CA |

**Figure S2. Secondary structure and conserved boxes of EcZraS-CD:** (A) Cartoon and stick diagram of H-box residues (deep salmon) in the DHp helix hα1 of EcZraS-CD. (B) Cartoon representation of the CA domain showing N (deep teal), G1 (green cyan), F (cyan), G2 (forest), G3 (gray50) boxes and the ATP lid (deep purple). Hydrophobic residues of gripper region (marine) and Phe of F-box, are also shown in stick representation. The ligand ADP (gray90) is shown as a stick and $Mg^{2+}$ (green) as a sphere. (C) and (E) Secondary structure of the DHp and CA domains. (D) Conserved boxes with their sequence stretch, conserved residues, and location in EcZraS-CD.

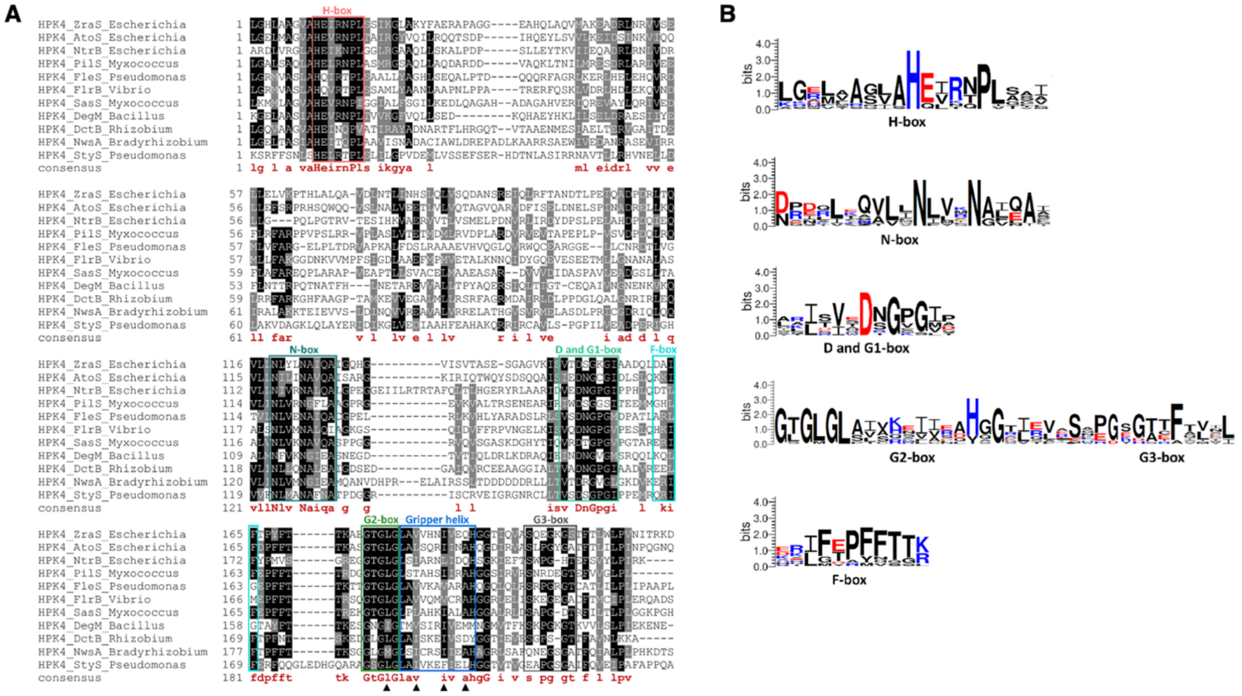

**Figure S3. Multiple sequence alignment and sequence consensus of conserved domains of EcZraS-CD:** (A) Sequence alignment showing conserved regions of kinase domains of sensor histidine kinases belonging to HPK4 family. Sequence consensus are represented below the alignment in maroon. Black triangles represent residues forming sticky fingers that help CA domain bind to the DHp bundle. (B) Logo depicting sequence consensus in H, N, F, G1, G2 and G3 boxes of HPK4 family representatives used in A.

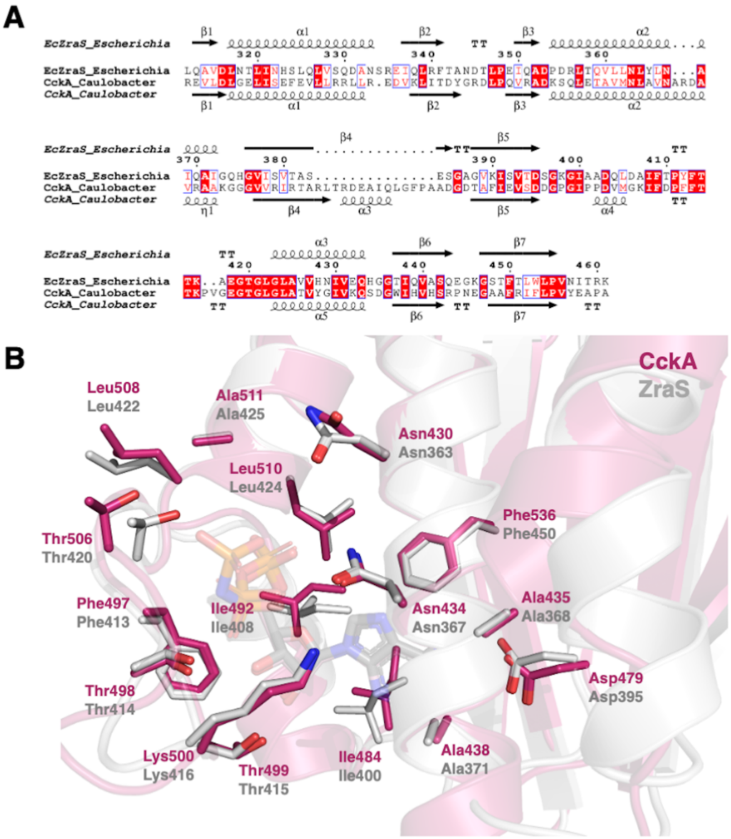


**Figure S4. Comparison of CA domain of EcZraS-CD and CckA:** (A) Structural alignment of ZraS with CckA. (B) Conserved residues enclosing the ligand binding pocket in the CA domain of ZraS (gray90) and CckA (hotpink).

**A**

r.m.s.d. values for EcZraS-CD chains

| Chains | cC | cD |
|---|---|---|
| cA | 1.022 | 3.070 |
| cB | 2.430 | 0.580 |

**B**

r.m.s.d. values for the DHp residue windows

| cA | 254-260 | 261-267 | 268-274 | 275-281 | 282-288 | 289-295 | 296-302 |
|---|---|---|---|---|---|---|---|
| cA | 0 | 0 | 0 | 0 | 0 | 0 | 0 |
| cB | 0.428 | 0.229 | 0.126 | 0.277 | 0.200 | 0.151 | 0.193 |
| cC | 0.293 | 0.188 | - | - | 0.228 | 0.099 | 0.141 |
| cD | 0.718 | 0.200 | 0.187 | 0.418 | 0.198 | 0.144 | 0.155 |

**C**

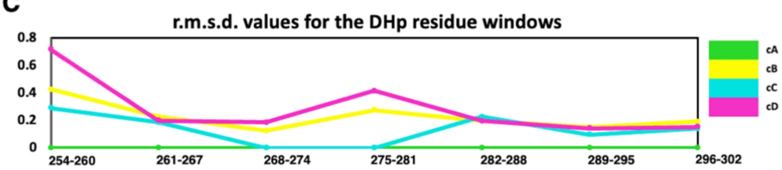


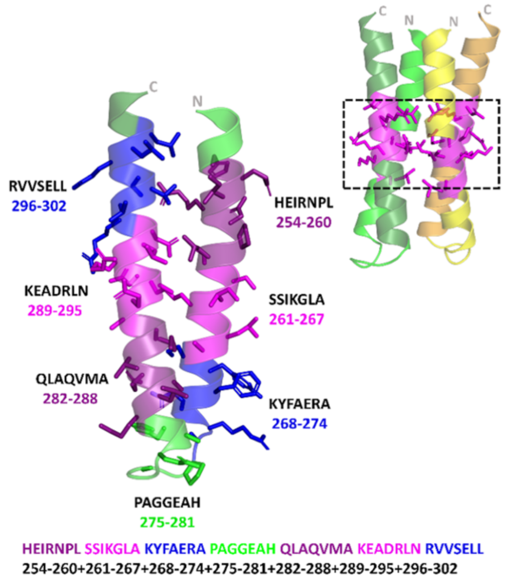


**Figure S5. Root mean square deviation (r.m.s.d.) (Å) values for the chains of EcZraS-CD:** (A) R.m.s.d. (Å) values calculated using superpose (PyMol) for the chains of dimers AB and CD. (B) R.m.s.d. (Å) values calculated using Pair_fit algorithm (PyMol) for the 7 residue windows of DHp domains in dimer AB and CD. Chain cA is used a reference for comparison. (C) Graphical representation of the r.m.s.d. (Å) values obtained in using Pair_fit. Pair_fit values for the region 268-274 and 275-281 in cC could not be obtained due to chain break.

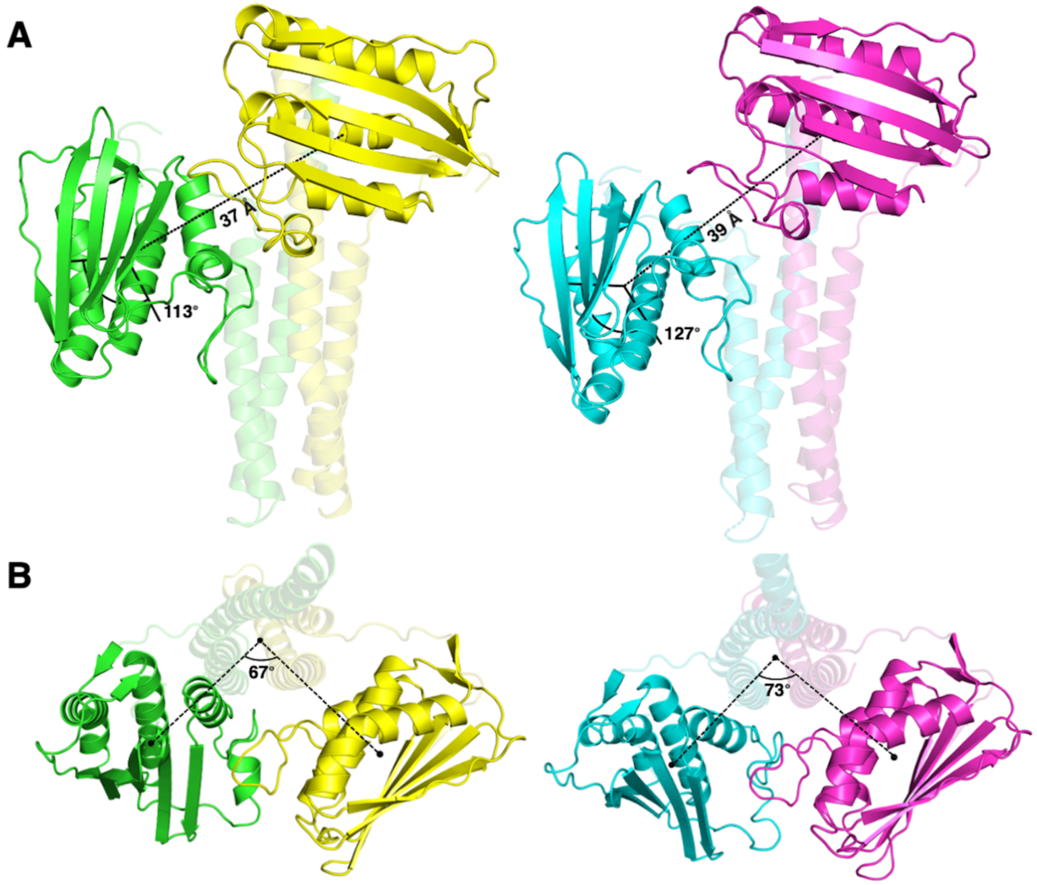


**Figure S6. Angles (°) and distances (Å) between CA domains of EcZraS-CD:** (A) Angle and distance between CA domains in dimers AB and CD calculated using angle_between_domains (PyMol) (33). (B) Angle of separation measured using centroids of the CA domains in dimers AB and CD shown in top view (33). Change in the orientation of the CA domain facilitates angular changes without affecting the visible separation between the domains.

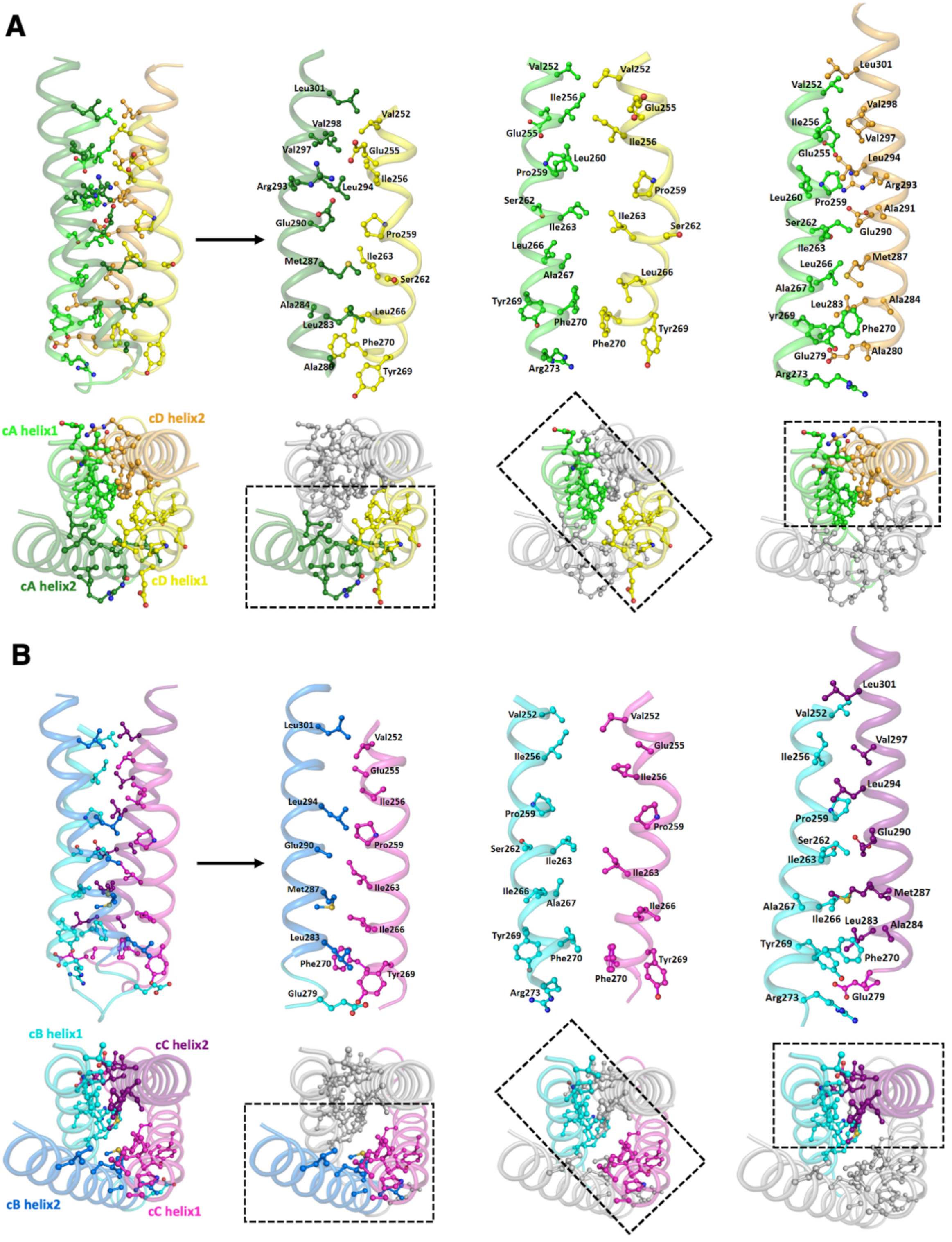


**Figure S7. Interactions at the interface of different DHp bundle helices of dimers AB and CD.** (A) Ball and stick representation of residues interacting at the DHp interface of helices in dimer AB. (B) Ball and stick representation of residues interacting at the DHp interface of helices in dimer CD.

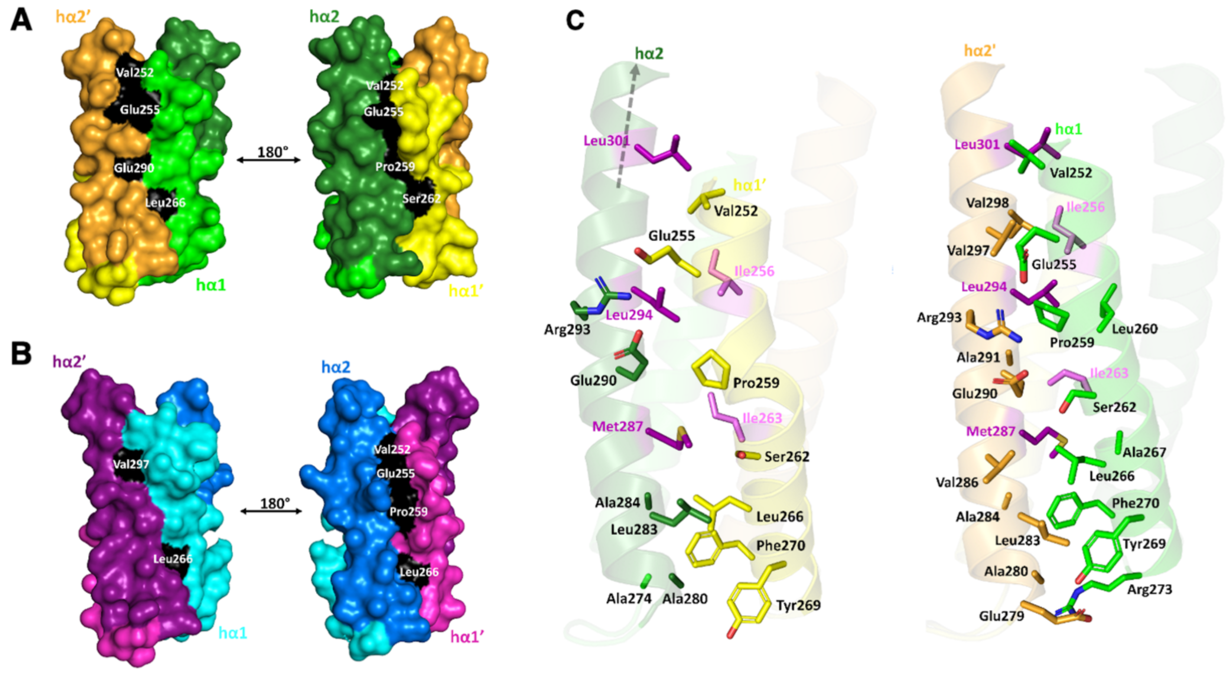


**Figure S8. Variable helical interface interactions of EcZraS-CD DHp domain** (A) and (B) “c” and “g” residues of DHp hα1-hα2’ and hα1’-hα2 of dimer AB and CD respectively, interacting directly with the inner core forming “d” residues, aiding in reorganization of coil layers. (C) Differential residue interactions along the DHp bundle interface between helices hα1’-hα2 and hα1-hα2’ in dimer AB.

**Table S1. Crick angle deviation values per layer of EcZraS-CD DHp (°):** Tabular representation of average Crick angle deviation (°) values for the residue layers, calculated using SamCC-Turbo (42) for the dimers AB and CD. "Layers" are the residue layers predicted according to SamCC-Turbo and "ad Layers" include the identified layers with a and d residues.

| ad Layers | Layers | Dimer AB | Dimer CD |
|---|---|---|---|
| | 0 | 8.13 | 6.45 |
| | 1 | 7.99 | 7.56 |
| adL2 | 2 | 5.99 | 5.58 |
| | 3 | 6.94 | 7.6 |
| | 4 | 6.87 | 5.2 |
| | 5 | 5.92 | 3.34 |
| adL3 | 6 | 7.99 | 4.3 |
| | 7 | 5.6 | 3.18 |
| | 8 | 6.25 | 4.62 |
| adL4 | 9 | 7.16 | 4.99 |
| | 10 | 7.99 | 6.58 |
| | 11 | 5.87 | 5.77 |
| | 12 | 6.3 | 7.66 |
| adL5 | 13 | 8.02 | 10.96 |
| | 14 | 8.47 | 12.03 |
| | 15 | 8.47 | 12.05 |

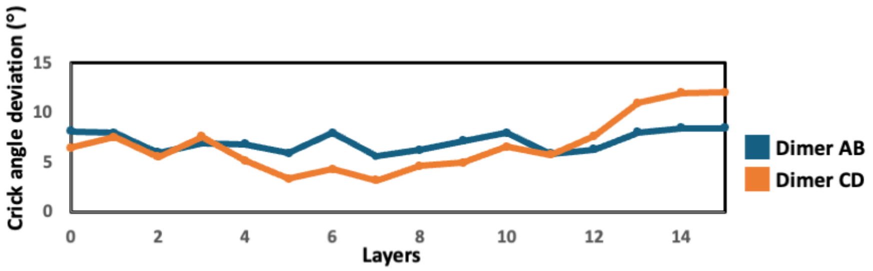


**Figure S8. Crick angle deviation (°) values per layer:** Line graph representation of the data in table S1. The variation in Crick angle deviation values (°) suggest reorganisation of the DHp core packing upon rotation of constituent helices in the bundle.

**Table S2. Crick angle deviation (°) values per residue of EcZraS-CD DHp helix-1 (254-269):** Tabular representation of the Crick angle deviation (°) values of the residues in the DHp helix-1 calculated using SamCC-Turbo (42).

| Residue no. | cA hα1 | cC hα1 | cB hα1' | cD hα1' |
|---|---|---|---|---|
| **254** | 4.36 | 1.98 | 33.72 | 37.62 |
| **255** | 3.1 | 7.14 | 30.37 | 35.93 |
| **256** | 5.19 | 12.73 | 23.27 | 21.68 |
| **257** | 10.51 | 16.27 | 18.7 | 22.32 |
| **258** | 8.77 | 10.58 | 16.73 | 11.23 |
| **259** | 4.06 | 6.17 | 12.46 | 4.07 |
| **260** | 6.32 | 9.06 | 20.25 | 4.39 |
| **261** | 7.6 | 7 | 10.02 | 5.02 |
| **262** | 10.05 | 11.55 | 10.61 | 6.12 |
| **263** | 12.07 | 12.2 | 12.39 | 11.06 |
| **264** | 14.51 | 11.84 | 15.08 | 16.05 |
| **265** | 9.58 | 9.97 | 15.04 | 14.12 |
| **266** | 8.6 | 13.12 | 17.73 | 14.17 |
| **267** | 10.74 | 19.33 | 24.16 | 18.94 |
| **268** | 15.44 | 21.63 | 23.32 | 23.54 |
| **269** | 19.24 | 23.77 | 22.82 | 24.27 |

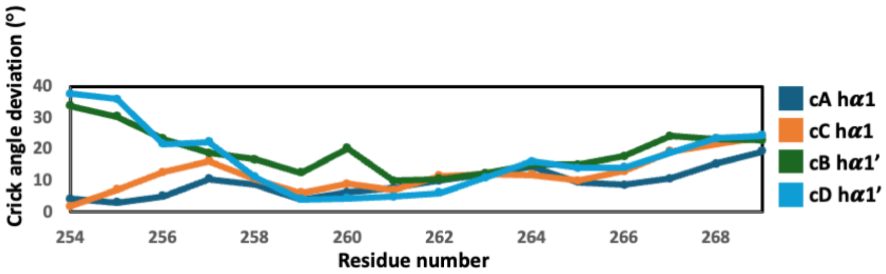


**Figure S9. Plot of Crick angle deviation (°) values per residue (254-269):** Graphical representation of the data present in the Table S2. The variation of Crick angle deviation (°) per residue is larger towards the N-termini, and minimal towards the C-termini.

**Table S3. Crick angle deviation (°) values per residue of EcZraS-CD DHp helix-2 (285-300):** Tabular representation of the Crick angle deviation (°) values of the residues in the DHp helix-2 calculated using SamCC-Turbo (42).

| Residue no. | cA hα2 | cC hα2 | cB hα2' | cD hα2' |
|---|---|---|---|---|
| **285** | -11.76 | 0.71 | 3.57 | -0.54 |
| **286** | -10.87 | 1.49 | 5.99 | 1.45 |
| **287** | -10.08 | 1.92 | 7.26 | 3.63 |
| **288** | -8.91 | 0.65 | 7.79 | 2.72 |
| **289** | -8.17 | -1.08 | 7.03 | 0.08 |
| **290** | -7.4 | -1.14 | 9.78 | -0.43 |
| **291** | -5.41 | -1.53 | 9.58 | -1.76 |
| **292** | -4.68 | 1.56 | 9.03 | -0.74 |
| **293** | -4.13 | 1.19 | 8.92 | -0.49 |
| **294** | -4.23 | 1.58 | 9.63 | 2.17 |
| **295** | -4.89 | 1.04 | 12.05 | 2.07 |
| **296** | -7.41 | -2.56 | 9.39 | 1.56 |
| **297** | -9.16 | -7.58 | 7.73 | -0.6 |
| **298** | -11.32 | -10.09 | 6.81 | -2.01 |
| **299** | -11.87 | -10.7 | 10.35 | -2.13 |
| **300** | -15.37 | -12.91 | 9.82 | -0.9 |

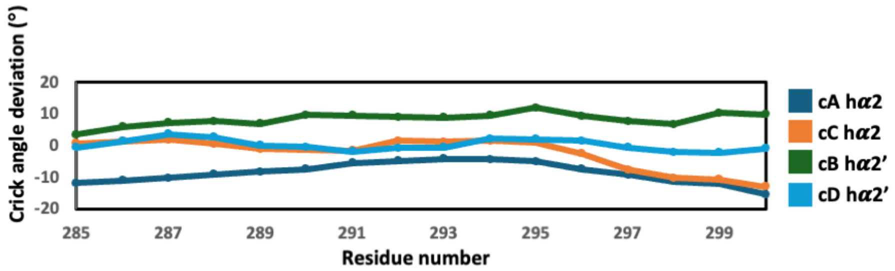


**Figure S10. Plot of Crick angle deviation (°) values per residue (285-300):** Graphical representation of the data present in the Table S3. The displacement of Crick angle deviation (°) per residue in cA and cB is relatively large as compared to that of cC and cD.

**Table S4. Axial shift (Å) values per residue of EcZraS-CD DHp helix-1 (254-269):** Tabular representation of the axial shift values (Å) of the residues in the DHp helix-1 calculated using SamCC-Turbo (42).

| Residue no. | cA hα1 | cC hα1 | cB hα1' | cD hα1' |
|---|---|---|---|---|
| **254** | 0.44 | -0.57 | -3.08 | 3.03 |
| **255** | 0.47 | -0.43 | 3.11 | 3.07 |
| **256** | 0.35 | 0.08 | 3.23 | 2.68 |
| **257** | 0.29 | -0.38 | 3.59 | 3.85 |
| **258** | 0.49 | -0.05 | 3.83 | 3.69 |
| **259** | 0.79 | 0.54 | 3.33 | 2.92 |
| **260** | 0.55 | 0.32 | 3.81 | 3.14 |
| **261** | 0.73 | 0.1 | 3.32 | 3.18 |
| **262** | 0.73 | 0.32 | 3.27 | 2.9 |
| **263** | 0.8 | 0.19 | 3.13 | 2.84 |
| **264** | 0.92 | -0.21 | 3 | 3.16 |
| **265** | 0.7 | -0.08 | 3.11 | 2.94 |
| **266** | 0.83 | -0.01 | 3.03 | 2.86 |
| **267** | 0.69 | 0.19 | 3.19 | 2.67 |
| **268** | 0.79 | -0.18 | 3.07 | 2.83 |
| **269** | 0.67 | -0.03 | 3.16 | 2.87 |

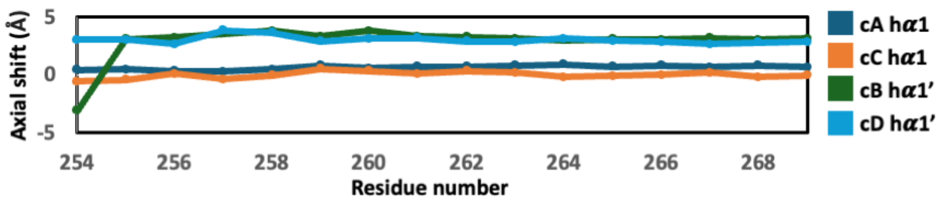


**Figure S11. Plot of Axial shift (Å) values per residue (254-269):** Graphical representation of the data present in the Table S4. The plot shows the axial shift (Å) per residue observed in cA and cB of dimer AB are compared to that of the corresponding chains cC and cD of dimer CD.

**Table S5. Axial shift (Å) values per residue of EcZraS-CD DHp helix-2 (285-300):** Tabular representation of the axial shift (Å) values of the residues in the DHp helix-2 calculated using SamCC-Turbo (42).

| Residue no. | cA hα2 | cC hα2 | cB hα2' | cD hα2' |
|---|---|---|---|---|
| **285** | 2.8 | -1.26 | -1.03 | -1.59 |
| **286** | -2.49 | -0.91 | -1.38 | -1.74 |
| **287** | -2.4 | -0.95 | -1.47 | -1.91 |
| **288** | -2.44 | -1.25 | -1.42 | -1.6 |
| **289** | -2.28 | -1.08 | -1.52 | -1.79 |
| **290** | -2.19 | -0.71 | -1.73 | -2.24 |
| **291** | -2.32 | -0.66 | -1.61 | -2.37 |
| **292** | -2.3 | -0.94 | -1.71 | -2.28 |
| **293** | -2.25 | -1.16 | -1.8 | -2.12 |
| **294** | -2.66 | -0.81 | -1.7 | -2.64 |
| **295** | -2.27 | -1.5 | -1.84 | -1.96 |
| **296** | -2.75 | -1.55 | -1.57 | -2.09 |
| **297** | -2.38 | -1.23 | -1.5 | -2.23 |
| **298** | -1.74 | -0.4 | -1.84 | -2.36 |
| **299** | -1.75 | -0.75 | -1.83 | -1.89 |
| **300** | -1.63 | -0.62 | -1.88 | -1.84 |

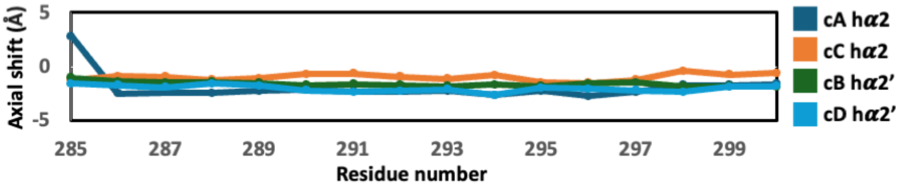


**Figure S12. Plot of Axial shift (Å) values per residue (285-300):** Graphical representation of the data present in the Table S5.

**Table S6: Axial radius (Å) values per residue of EcZraS-CD DHp helix-1 (254-269):** Tabular representation of the axial radius (Å) values of the residues in the DHp helix-1 calculated using SamCC-Turbo (42).

| Residue no. | cA hα1 | cC hα1 | cB hα1' | cD hα1' |
|---|---|---|---|---|
| **254** | 6.23 | 7.02 | 7.85 | 8.22 |
| **255** | 6.22 | 7.2 | 7.86 | 8.34 |
| **256** | 6.11 | 7.18 | 7.75 | 8.22 |
| **257** | 6.15 | 7.31 | 7.91 | 8.57 |
| **258** | 5.88 | 6.98 | 7.95 | 8.3 |
| **259** | 5.98 | 6.91 | 8.03 | 8.03 |
| **260** | 5.98 | 6.79 | 8.19 | 7.72 |
| **261** | 5.86 | 6.59 | 7.84 | 7.39 |
| **262** | 5.76 | 6.43 | 7.64 | 7.1 |
| **263** | 5.66 | 6.37 | 7.4 | 6.93 |
| **264** | 5.65 | 6.22 | 7.24 | 6.83 |
| **265** | 5.58 | 6.09 | 7.05 | 6.67 |
| **266** | 5.58 | 6.16 | 6.96 | 6.64 |
| **267** | 5.6 | 6.25 | 6.9 | 6.67 |
| **268** | 5.66 | 6.28 | 6.83 | 6.66 |
| **269** | 5.77 | 6.26 | 6.83 | 6.68 |

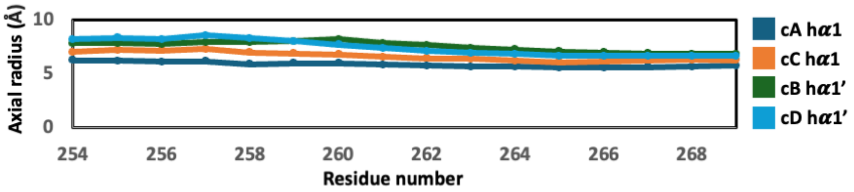


**Figure S13. Plot of Axial radius (Å) values per residue (254-269):** Graphical representation of the data present in the Table S6.

**Table S7. Axial radius (Å) values per residue of EcZraS-CD DHp helix-2 (285-300):** Tabular representation of the axial radius (Å) values of the residues in the DHp helix-2 measured using SamCC-Turbo (42).

| Residue no. | cA hα2 | cC hα2 | cB hα2' | cD hα2' |
|---|---|---|---|---|
| **285** | 8.38 | 8.11 | 7.69 | 7.74 |
| **286** | 8.28 | 7.95 | 7.52 | 7.57 |
| **287** | 8.23 | 7.82 | 7.36 | 7.43 |
| **288** | 8.19 | 7.82 | 7.29 | 7.36 |
| **289** | 8.15 | 7.71 | 7.23 | 7.3 |
| **290** | 8.18 | 7.68 | 7.23 | 7.28 |
| **291** | 8.2 | 7.69 | 7.2 | 7.19 |
| **292** | 8.28 | 7.73 | 7.23 | 7.11 |
| **293** | 8.36 | 7.7 | 7.27 | 7.04 |
| **294** | 8.44 | 7.68 | 7.33 | 7.11 |
| **295** | 8.33 | 7.76 | 7.42 | 7.14 |
| **296** | 8.29 | 7.7 | 7.45 | 7.36 |
| **297** | 8.16 | 7.65 | 7.5 | 7.39 |
| **298** | 8.08 | 7.57 | 7.53 | 7.38 |
| **299** | 8.08 | 7.68 | 7.53 | 7.35 |
| **300** | 7.98 | 7.71 | 7.51 | 7.41 |

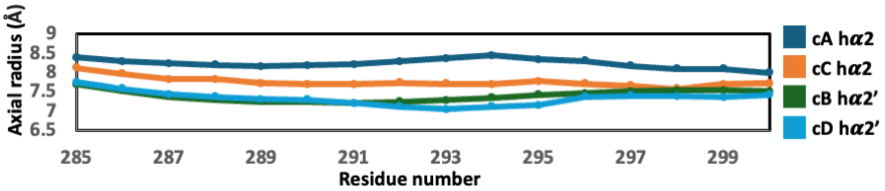


**Figure S14. Plot of Axial radius (Å) values per residue (284-300):** Graphical representation of the data present in the Table S7.

**Table S8. Prediction of heptad repeats in linker and DHp helix-1 of EcZraS-CD:** Tabular representation of the heptad positions predicted using the program TWISTER (41). The region predicted to be the stutter is highlighted in yellow.

| Residue no. | Residue | Dimer AB | Dimer CD |
|---|---|---|---|
| 226 | L | | f |
| 227 | R | | g |
| 228 | S | | a |
| 229 | R | | b |
| 230 | Q | c | c |
| 231 | L | d | a |
| 232 | L | e | b |
| 233 | Q | f | c |
| 234 | D | g | d |
| 235 | E | a | e |
| 236 | M | b | f |
| 237 | K | c | g |
| 238 | R | d | d |
| 239 | K | e | e |
| 240 | E | f | f |
| 241 | K | g | g |
| 242 | L | a | a |
| 243 | V | b | b |
| 244 | A | c | c |
| 245 | L | d | d |
| 246 | G | e | e |
| 247 | H | f | f |
| 248 | L | g | g |
| 249 | A | a | a |
| 250 | A | b | b |
| 251 | G | c | c |
| 252 | V | z | z |
| 253 | A | a | a |
| 254 | H | b | b |
| 255 | E | c | c |
| 256 | I | d | d |
| 257 | R | e | e |
| 258 | N | f | f |
| 259 | P | g | g |
| 260 | L | a | a |
| 261 | S | b | b |
| 262 | S | c | c |
| 263 | I | d | d |
| 264 | K | e | e |
| 265 | G | f | f |
| 266 | L | g | g |
| 267 | A | a | a |
| 268 | K | b | b |
| 269 | Y | c | c |
| 270 | F | d | d |
